# A transcriptome atlas of leg muscles from healthy human volunteers reveals molecular and cellular signatures associated with muscle location

**DOI:** 10.1101/2022.06.01.494335

**Authors:** Tooba Abbassi-Daloii, Salma el Abdellaoui, Lenard M. Voortman, Thom Veeger, Davy Cats, Hailiang Mei, Duncan E. Meuffels, Ewoud van Arkel, Peter A.C ’t Hoen, Hermien E. Kan, Vered Raz

**Affiliations:** Department of Human Genetics, Leiden University Medical Center, Leiden, 2333 ZA, the Netherlands; Division of Cell and Chemical Biology, Leiden University Medical Centre, Leiden, 2333 ZA, the Netherlands; C.J. Gorter Center for High Field MRI, Department of Radiology, Leiden University Medical Center, Leiden, 2333 ZA, the Netherlands; Sequencing Analysis Support Core, Leiden University Medical Center; Orthopedic and Sport Medicine Department, Erasmus MC, University Medical Center Rotterdam, 3015 GD, Rotterdam, the Netherlands; Orthopedics, Medisch Centrum Haaglanden, 2501 CK, Den Haag, the Netherlands; Centre for Molecular and Biomolecular Informatics, Radboud Institute for Molecular Life Sciences, Radboud University Medical Center, 6525 EX, Nijmegen, the Netherlands; Duchenne Center Netherlands, 5591 VE, Heeze, the Netherlands

**Keywords:** Skeletal muscles, mRNA expression atlas, molecular and cellular signatures, Capillary density, Myofiber type, Homeobox genes

## Abstract

Skeletal muscles support the stability and mobility of the skeleton but differ in biomechanical properties and physiological functions. The intrinsic factors that regulate muscle-specific characteristics are poorly understood. To study these, we constructed a large atlas of RNA-seq profiles from six leg muscles and two locations from one muscle, using biopsies from 20 healthy young males. We identified differential expression patterns and cellular composition across the seven tissues using three bioinformatics approaches confirmed by large-scale newly developed quantitative immune-histology procedures. With all three procedures, the muscle samples clustered into three groups congruent with their anatomical location. Concomitant with genes marking oxidative metabolism, genes marking fast- or slow-twitch myofibers differed between the three groups. The groups of muscles with higher expression of slow-twitch genes were enriched in endothelial cells and showed higher capillary content. In addition, expression profiles of Homeobox (*HOX*) transcription factors differed between the three groups and were confirmed by spatial RNA hybridization. We created an open-source graphical interface to explore and visualize the leg muscle atlas (https://tabbassidaloii.shinyapps.io/muscleAtlasShinyApp/). Our study reveals molecular specialization of human leg muscles and provides a novel resource to study muscle-specific molecular features, which could be linked with (patho)physiological processes.

## Introduction

Skeletal muscles have *grosso modo* similar functions, generate the force for mobility and skeleton support, and maintain the body homeostasis. However, skeletal muscles differ in biomechanical and physiological features. These features include the size and contractile properties of the motor units and myofibers, differences in shortening velocity, resistance to fatigue, and differences in innervation and perfusion (Valentine 2017). Yet, the molecular and cellular differences that contribute to this muscle specialization are not fully understood. A molecular atlas for different skeletal muscles could assist in deciphering the molecular basis of muscle-specific physiological features. Such an atlas may also be used to study differential muscle involvement in various conditions, such as muscular dystrophies, myopathies, differences in regenerative potential, physiological compensation in sports and sarcopenia. In aging and muscle diseases, Muscle involvement it has been observed that some muscles were affected at an earlier age than others, and that this muscle involvement pattern can be characteristic for a given disease (Carlier, Laforet et al. 2011, Raz, Henseler et al. 2015, Albayda, Christopher-Stine et al. 2018, Brogna, Cristiano et al. 2018, Diaz-Manera, Fernandez-Torron et al. 2018, Servian-Morilla, Cabrera-Serrano et al. 2020). Several studies suggested that muscle-specific intrinsic molecular factors may explain this muscle involvement pattern (Kang, Kho et al. 2005, Rahimov, King et al. 2012, Huovinen, Penttila et al. 2015, Raz, Henseler et al. 2016, Terry, Zhang et al. 2018, Hettige, Tahir et al. 2020, Xi, Langerman et al. 2020). For example, differences in the cellular pathways and myofiber type (slow- and fast-twitch myofibers) composition between muscles could play a role (De Micheli, Spector et al. 2020, Rubenstein, Smith et al. 2020, Xi, Langerman et al. 2020), but may not fully explain the muscle involvement patterns.

Most of the studies characterizing the molecular variation between muscles were performed in mice (Campbell, Gordon et al. 2001, Porter, Khanna et al. 2001, Haslett, Kang et al. 2005, von der Hagen, Laval et al. 2005, Raz, Riaz et al. 2018, Terry, Zhang et al. 2018, Hettige, Tahir et al. 2020), where muscle-specific mRNA profiles were linked to distinct myofiber type composition (Campbell, Gordon et al. 2001, Raz, Riaz et al. 2018, Hettige, Tahir et al. 2020). Since human muscle-related pathologies are not always recapitulated in mouse models (van Putten, Lloyd et al. 2020), understanding molecular variations between skeletal muscles should be performed in human samples. Only a few studies compared mRNA profiles between muscles from healthy human adults, and these studies face several limitations. Skeletal muscles are highly affected by age (McCormick and Vasilaki 2018, Aversa, Zhang et al. 2019), yet, the age range in previous studies was broad (Kang, Kho et al. 2005, Huovinen, Penttila et al. 2015). Moreover, the numbers of sampled muscles and subjects were limited (Abbassi-Daloii, Kan et al. 2020). A study using postmortem material (Kang, Kho et al. 2005) only partly reflects molecular composition in living muscles due to storage in cooling conditions. Understanding muscle involvement in different pathologies can benefit from a molecular atlas of human muscles.

We generated a transcriptome atlas from six leg muscles and two locations from one muscle to explore molecular variations within and between muscles. Paired samples were obtained from 20 healthy male subjects of 25 ± 3.6 years old. We show that the seven muscle tissues clustered into three groups, distinguished by cell type composition and mRNA expression profiles. We confirmed the transcriptome analyses with large-scale quantitative immunohistochemistry and RNA *in situ* hybridization procedures. We discuss the value of this skeletal muscle atlas resource to understand human health and pathologies affecting skeletal muscle tissues.

## Results

### Transcriptome atlas of adult human skeletal muscles

To determine molecular signatures marking leg muscles, we generated a transcriptome atlas of human skeletal muscles by sequencing biopsies from five upper leg muscles, gracilis (GR), semitendinosus (ST), rectus femoris (RF), vastus lateralis (VL), and vastus medialis (VM) muscles, and one lower leg muscle, gastrocnemius lateralis (GL) (Figure 1A). We also investigated molecular differences within one muscle by including biopsies from the middle and distal end of the semitendinosus muscle (STM and STD, respectively). These two biopsies were treated as independent muscle samples in subsequent analyses (Figure 1B and C). In total, 128 samples from 20 individuals (aged 25 ± 3.6 yr) were analyzed (Supplementary Figure S1), making this currently the largest freely available human muscle atlas. Supplementary Table S1 shows the sample characteristics.

**Figure 1.**
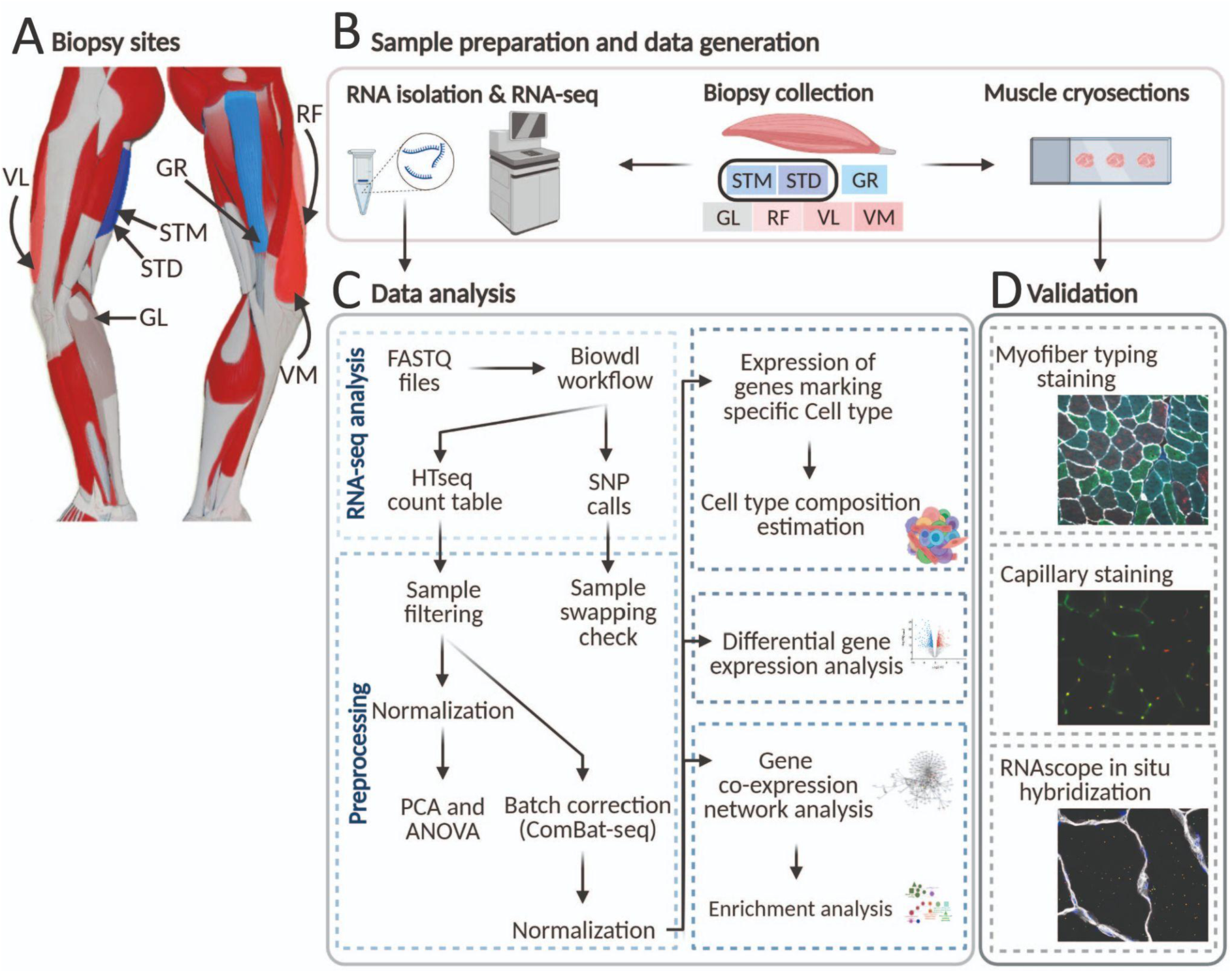
An overview of biopsies’ location and the study workflow. A) A schematic overview of the leg muscles. Arrows point to the muscles that were included in this study. The biopsies, with exception of STM (semitendinosus-middle), were taken from the distal area. B-D) The study overview includes cryosectioning, RNA-isolation and sequencing (B) data analysis (C) and validations (D).

### Variation in cell type composition between different muscles

Skeletal muscle is a heterogeneous tissue containing multiple cell types. The differences in the abundance of these cell types can be reflected in bulk RNA-seq profiles. Therefore, we used RNA-seq data to first explore possible cell type heterogeneity between leg muscles. We summarized the expression level of genes marking each cell type present in human skeletal muscles by calculating their first principal components (eigenvectors) (Supplementary Table S2). We used the eigenvalues of the eigenvectors representing the different cell types to cluster the muscles (Figure 2A) and to identify cell types with significant differences in relative abundance between muscles (Supplementary Figure S2A). The muscle tissues clustered into three groups, Group 1 (G1): GR, STM, and STD; Group 2 (G2): RF, VL, and VM; GL was the only muscle in Group 3 (G3) (Figure 2A). The relative abundance of endothelial cells was statistically the most different between muscles, with higher abundance in G2 and G3 than in G1 (Figure 2A-B, Supplementary Figure S2A). Other cell types marking blood vessels, namely pericytes, post-capillary venule (PCV) endothelial cells, natural killer (NK) cells, T and B cells, and myeloid cells, clustered together with the endothelial cells and all showed a higher abundance in G2 and G3 compared with G1 (Figure 2A-B). These results could suggest a higher capillary density and blood perfusion in/of the muscles in G2 and G3.

**Figure 2.**
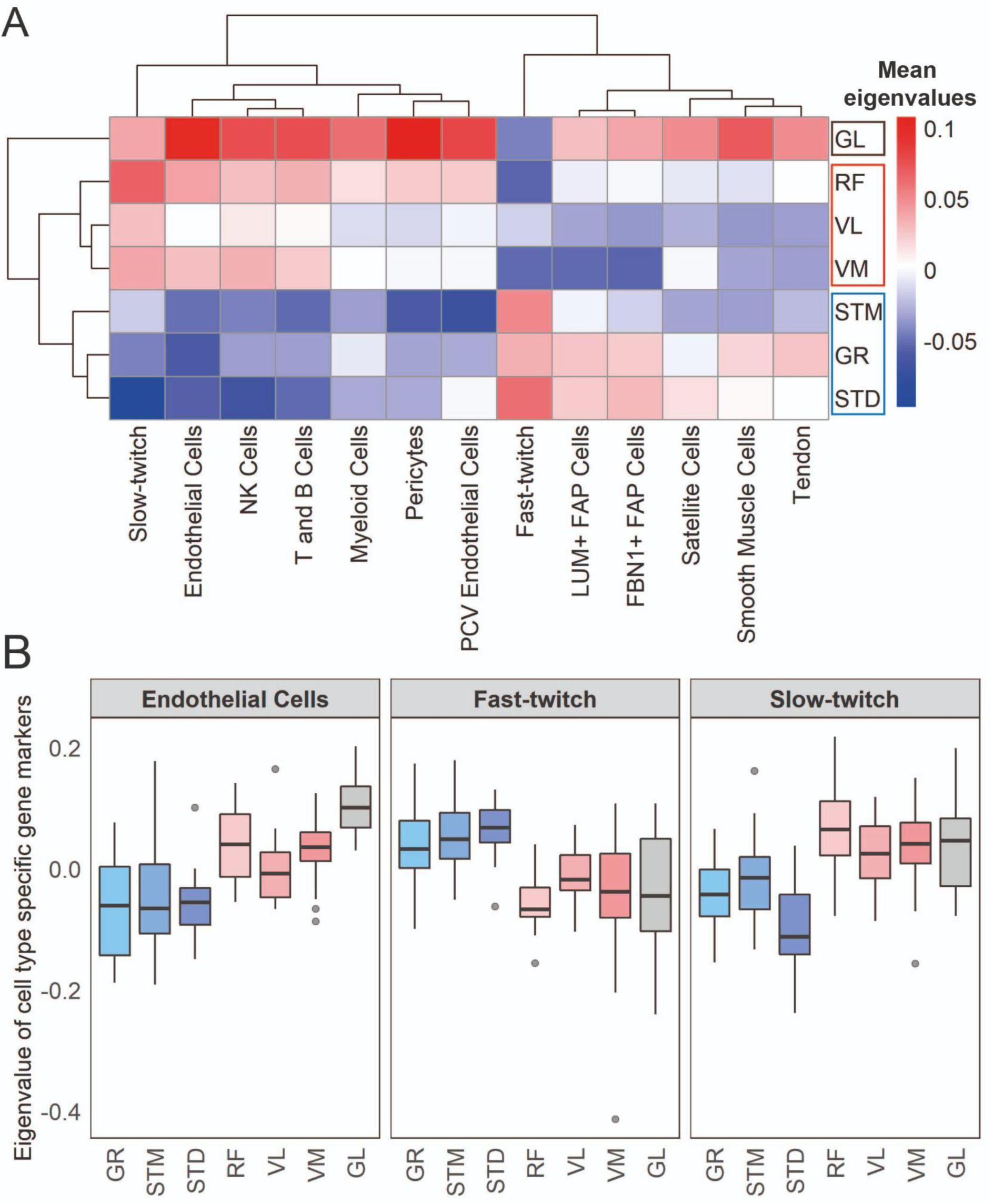
Muscles cluster into three main groups based on cell type composition. **A)** The heatmap shows the mean eigenvalues of genes marking each cell type across all the individuals. Each row shows a muscle, and each column shows a cell type. FAB stands for fibro-adipogenic progenitors. **B)** The boxplot shows the eigenvalues for the endothelial cells, fast-twitch, and slow-twitch myofibers per muscle. The boxes reflect the median and interquartile range.

Genes marking fast-twitch myofibers showed overall higher expression levels in G1, while slow-twitch genes were higher in G2 and G3 (Figure 2A-B). Differences in the relative abundance of non-muscle cell types, pericytes, immune cells, and endothelial cells, distinguished G3 from the G1 and G2 muscles (Supplementary Figure S2B, Figure 2A-B).

While STM and STD showed significant differences in the relative abundance of genes marking endothelial cells (higher expression in STD) and slow-twitch myofibers (higher expression in STM), there were no significant differences between ST and GR, and within the G2 muscles. This suggests that differences between regions of the same muscle may be larger than differences between distinct muscles (Supplementary Figure S2A).

### Further study of differences in myofiber type composition between groups of muscles

To confirm the differences in myofiber types between muscles, we performed immunofluorescence staining for all muscles with a mixture of antibodies to three MyHC isoforms and anti-laminin antibody (Figure 3A). We developed a semi-automated image processing workflow to segment the myofibers using laminin staining and to quantify the fluorescence intensity of each MyHC isoform per myofiber. We next identified myofiber types by clustering all the myofibers using the MFI values of the three MyHC isoforms. The vast majority of the myofibers (94%) were assigned to three major clusters (Figure 3B). Each myofiber cluster had a major MyHC isoform (Figure 3C). Consistent with our study in human *vastus lateralis* muscle (Raz, van den Akker et al. 2020), the results here suggest that the myofibers are generally not purely type -I, -IIA, or -IIX but contain a mix of myosin heavy chain isoforms. We observed relatively high correlations from 0.55 to 0.62 between normalized gene expression of the dominant MyHC in each cluster and the proportion of myofibers assigned to the corresponding cluster (Figure 3D-F). This correlation demonstrates the reliability of our RNA-seq-based assessment of MyHC expression.

**Figure 3.**
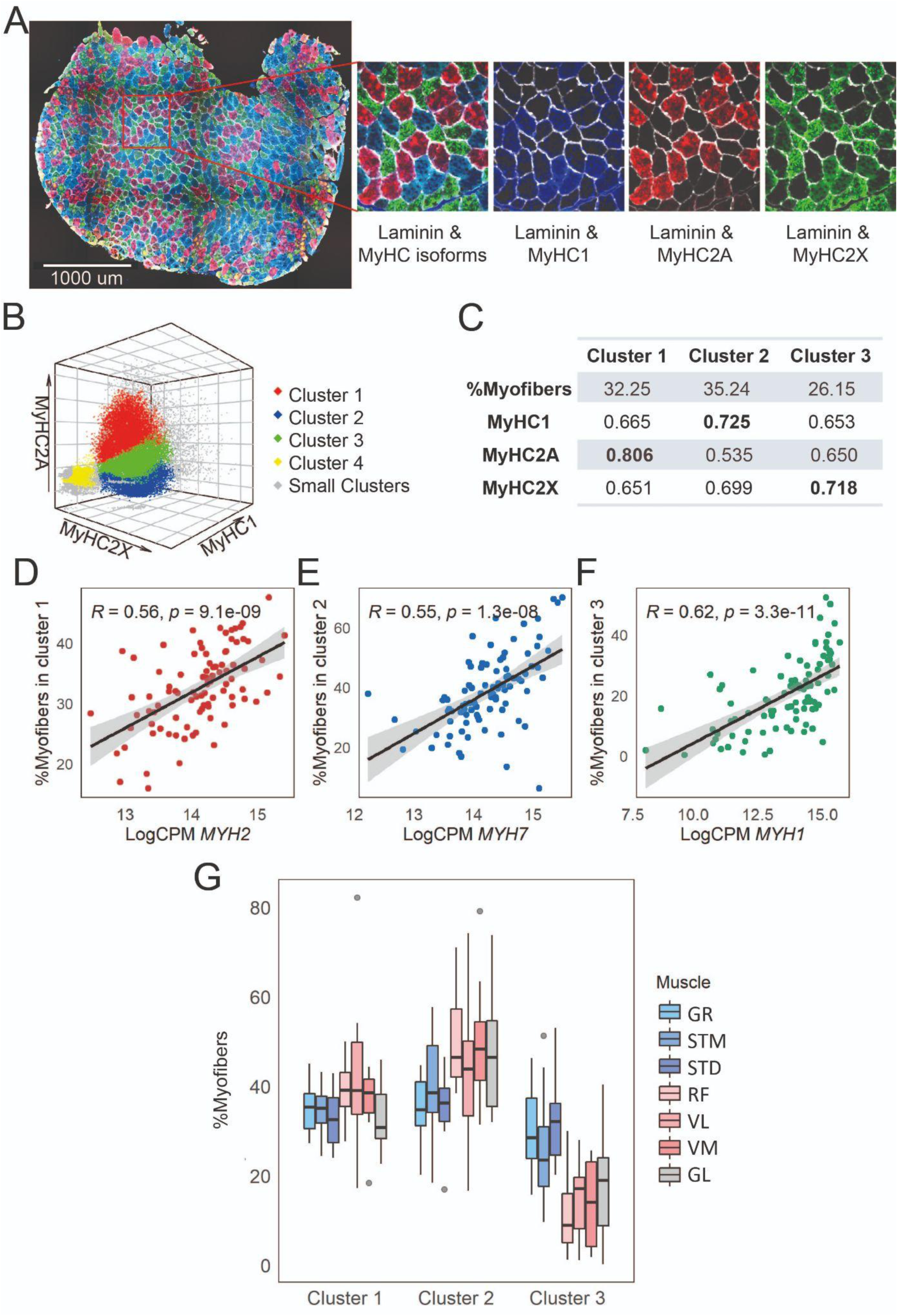
Myofiber type composition is consistent with the expression level of genes marking fast and slow-twitch myofibers. **A)** A representative immunostaining image. The overlay of each myosin heavy chain isoform and laminin are shown separately. **B)** The MFI of the three MyHC isoforms are plotted in 3-D. Each dot represents a myofiber. Myofibers in the three largest clusters are denoted with red (Cluster 1), blue (Cluster 2), and green (Cluster 3). The objects with low MFI values for all the isoforms are denoted in yellow (Cluster 4, ∼2% of all the dots). In gray are ∼4% of myofibers assigned to many small clusters. **C)** The table shows the proportion of myofibers assigned to each of the three largest clusters and the average MFI values for each isoform. **D-F)** Scatterplots show the proportion of the assigned myofibers to each of the largest clusters and the normalized expression of the gene coding the isoform with a relatively higher expression in that specific myofiber cluster. **G)** The boxplot shows the proportion of myofibers in the three largest clusters per muscle. Each muscle is depicted with a different color, with G1 muscles in blue, G2 muscles in red and the G3 muscle in grey. The boxes reflect the median and interquartile range.

In agreement with the results of the RNA-seq cell type composition analysis (Figure 2A-B), the quantitative histology analysis demonstrated a higher proportion of slow-twitch (oxidative) myofibers and a lower proportion of MyHC2X-dominated myofibers in G2 and GL (G3) muscles than in G1 muscles (Figure 3G, Supplementary Figure S3). The quantitative histology analysis further showed that G2 muscles had a higher proportion of MyHC2A-dominated myofibers than the G3 muscle (Figure 3G, Supplementary Figure S3), highlighting a distinct myofiber type composition of the GL muscle.

The myofiber composition results further showed a higher proportion of MyHC2X-dominated myofibers in STD than in STM, whereas STM had a higher proportion of MyHC1 and MyHC2A-dominated myofibers (Supplementary Figure S3). This confirms the existence of regional differences within a muscle (Bindellini, Voortman et al. 2021).

### Higher capillary density in GL

The RNA-seq cell type composition analysis suggested a higher proportion of endothelial and other cell types marking blood vessels in the GL muscle than in other muscles. To confirm this observation, we immunostained for the endothelial cells using antibodies against Endoglin (ENG) and CD31 proteins (Tey, Robertson et al. 2019). We included cryosections of GL and STM with the largest differences in the expression of genes marking endothelial cells (Figure 4A). We observed a higher proportion of CD31-positive areas in GL (Figure 4B), which was consistent with higher *CD31* RNA expression levels in this muscle (Figure 4C).

**Figure 4.**
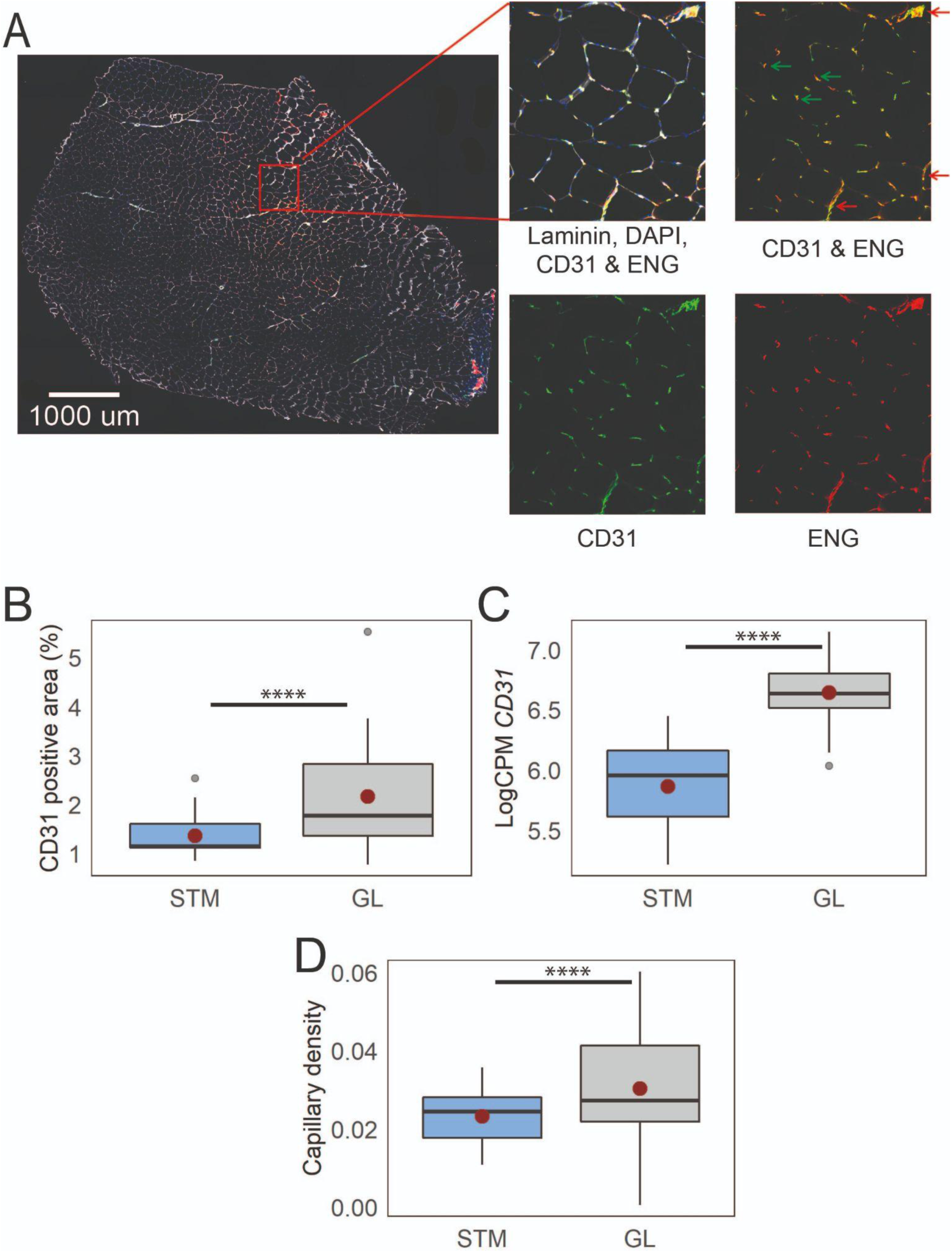
Immunostaining confirms higher capillary density in GL compared with STM muscles. **A)** A representative muscle cross section image immunostained with CD31, ENG, and laminin. An enlargement of the boxed region is shown on the right, with images of the three separate channels and an overlay. Examples of objects recognized as capillaries are shown by green arrows. Some examples of objects that were not considered as capillaries are shown by red arrows. **B)** The box plot shows the percentage of CD31 positive area in the two muscles. **C)** The box plot shows the normalized expression of *CD31* gene in the two muscles. **D)** The boxplot shows the estimated capillary density in the two muscles. The capillary density was defined as the number of objects (3-51 µm^2^) with an overlap between CD31 and ENG per unit cross-sectional area of the muscle. The boxes reflect the median and interquartile range (N = 19 per muscle). The red dots on the boxes show the mean. **** *P-value* < 1×10^-6^ (linear mixed-model).

Next, we determined the muscles’ capillary density by counting small circular objects stained positive for both CD31 and ENG (Wehrhan, Stockmann et al. 2011). We observed a higher capillary density in GL compared with STM (Figure 4D). This observation is consistent with a higher proportion of endothelial cell types in GL compared with muscles in G1 or G2.

### Gene expression profiles and molecular pathways distinguishing muscle clusters

We next investigated whether muscle-specific gene expression profiles, not explained by cell type composition, could also be found in our dataset. To this end, we determined the differentially expressed genes (DEGs) in every pairwise comparison (Supplementary Figure S4A, Supplementary Table S3). The DEGs that were driven by differences in cell type composition were excluded (Pearson’s R > 0.5 between gene expression levels and the eigenvector of any cell type). The proportion of DEGs that were not driven by cell type composition but discriminated each pair of muscles are shown in Figure 5A. The muscles clustered in a similar way as was observed in the cell type composition analysis: GR, STM, and STD (G1), RF, VL, and VM (G2), and GL (G3) (Figure 5A).

**Figure 5.**
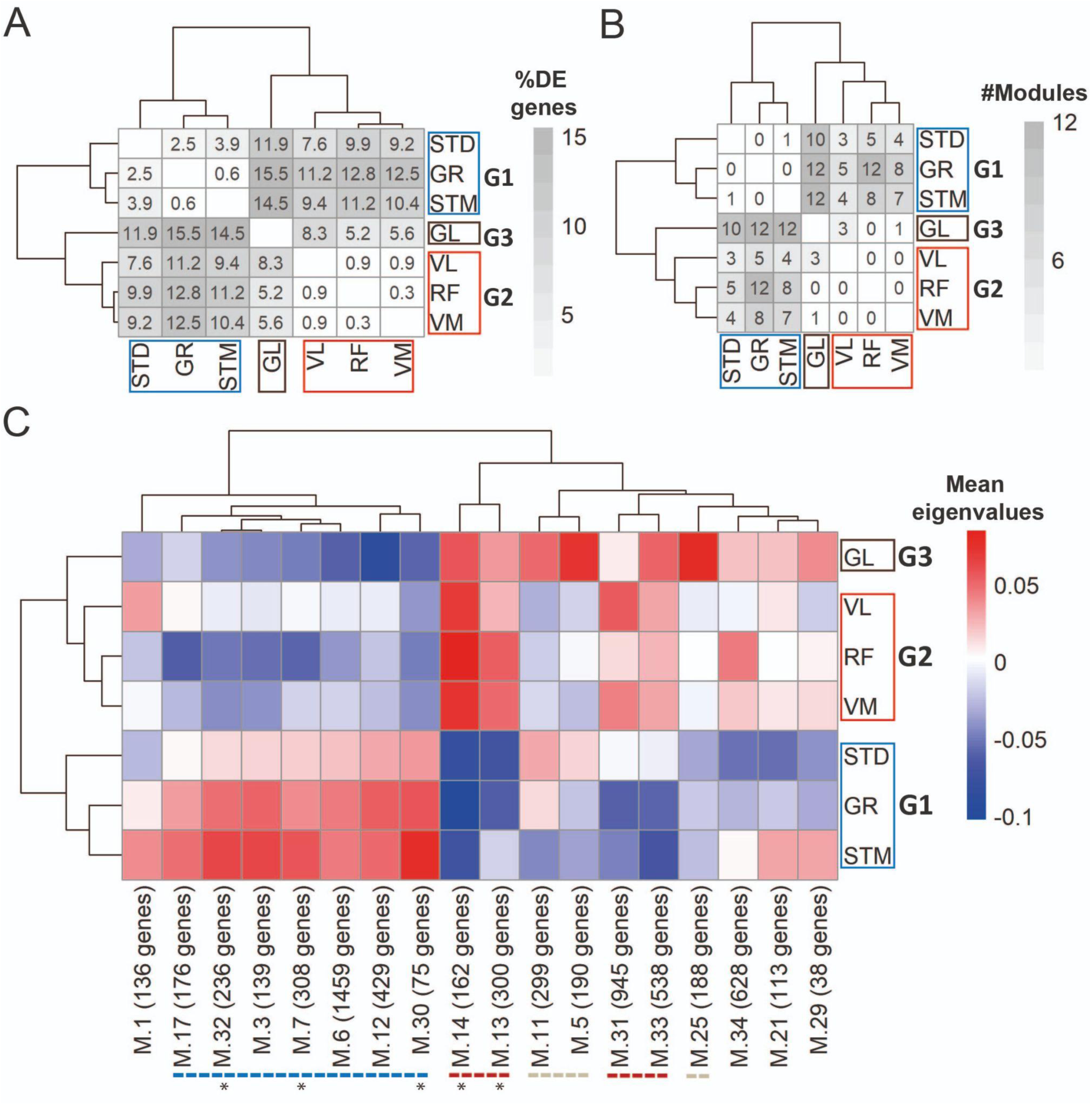
Gene expression differences between three groups of muscles not driven by cell type composition. **A)** Symmetric heatmap plot shows the percentage of DEGs in different pairwise comparisons. Genes with a high Pearson correlation (R > 0.5) with the eigenvector of any cell type are excluded. Each row or column represents a muscle. **B)** Symmetric heatmap plot shows the number of modules that were not driven by cell type composition and were significantly different in each pairwise comparison. Each row or column represents a muscle. **C)** The heatmap shows the modules that reflect the intrinsic differences between groups of muscles. Each row represents a muscle, and each column shows a muscle-related module that was not driven by cell type composition. Color-coded cells show the corresponding average of eigenvalues across all individuals (N = 20). Modules with an overall higher expression in G1 or G3 are underlined by a blue or gray dashed line, respectively. The red dashed line underlines the modules with an overall higher expression in both G2 and G3. The black asterisks show modules with the largest differences between the three groups of muscles.

To further study muscle-specific expression profiles, we applied weighted gene co-expression network analysis (WGCNA). We identified 35 modules of co-expressed genes (Supplementary Table S4). For each module, we calculated the module eigengene (ME) that represents gene expression levels of the genes in the module. We then implemented a pairwise comparison to find modules showing significant differences in every pairwise comparison (Supplementary Figure S4B-C). Out of the 35 modules, 27 showed a difference between at least two muscles (module size range: 38-1,459; containing 10,695 genes in total). Nine out of the 27 muscle-related modules had at least five genes marking a specific cell type and were therefore defined as modules driven by differences in cell type composition and were not considered for further analysis (Supplementary Figure S4D). Figure 5B shows the remaining modules that were not driven by cell type composition, nevertheless distinguished pairs of muscles. We then plotted the mean eigenvalues of muscle-related modules in a heatmap (Figure 5C) to determine the clustering of muscles based on the expression patterns of genes in the modules. The WGCNA-based clustering was consistent with the cell type composition and differential expression groups (Figure 5C). In total, seven out of the 18 muscle-related modules demonstrated higher expression levels in G1, four modules had higher expression in G2 and G3, and three modules demonstrated higher expression levels in G3 only (Figure 5C). In addition, although none of the modules showed distinct expression patterns between muscles in G2 and between ST and GR, M.21 module showed higher expression levels in STM than STD (Supplementary Figure S4C).

To explore the molecular and cellular pathways in the three groups, functional enrichment analysis was performed in the muscle-related modules. The most significantly enriched biological processes and molecular functions within these modules are listed in Table 1 (a complete list is in Supplementary Table S5).

**Table 1.** Top enrichment results of muscle-related modules not driven by cell type composition.

| Module | Term | FDR | #Enriched genes | Hub genes |
| --- | --- | --- | --- | --- |
| **Higher expression in G1 (GR, STM, and STD)** |  |  |  |  |
| M.30 (75 genes) | IRE1-mediated unfolded protein response | $4 \times 10^{-4}$ | 4 | |
| M.32 (236 genes) | Hormone-mediated signaling pathway | $9 \times 10^{-3}$ | 11 | *CARM1, WBP2, ZBTB7A* |
| M.7 (308 genes) | Negative regulation of nucleobase-containing compound metabolic process | $6 \times 10^{-4}$ | 51 | *CREBBP, DAB2IP, FOXK2, LARP1, RXRA, THRA* |
| | Chromatin organization | $1 \times 10^{-2}$ | 27 | *ARID1B, CREBBP, HUWE1* |
| M.17 (176 genes) | Chromatin modifying enzymes | $4 \times 10^{-3}$ | 11 | *HCFC1, SETD1A* |
| **Higher expression in G2 (RF, VL, and VM) and G3 (GL)** |  |  |  |  |
| M.31 (945 genes) | RNA splicing | $6 \times 10^{-5}$ | 54 | *SNRNP70* |
| | Histone modification | $3 \times 10^{-3}$ | 47 | *KAT2A* |
| M.33 (538 genes) | Apical junction complex | $2.4 \times 10^{-2}$ | 11 | *MICALL2* |
| M.13 (300 genes) | Mitochondrion | $4 \times 10^{-45}$ | 122 | *AIFM1, ATP5F1A, ATP5F1B, CKMT2, COQ9, DLD, DLST, FH, GHITM, HADHA, HADHB, IMMT, MFN2, NDUFS2, PDHA1, PDHB, TRAP1, UQCRC2* |
| M.14 (162 genes) | Anterior/posterior pattern specification | $4 \times 10^{-3}$ | 7 | *HOXA11* |
| **Higher expression in G3 (GL)** |  |  |  |  |
| M.5 (190 genes) | Regulation of lipid metabolic process | $8 \times 10^{-4}$ | 15 | *ADIPOQ, ADRA2A, CIDEA, LEP, LGALS12, PDE3B, SCD* |
| M.25 (188 genes) | Ameboidal-type cell migration | $5 \times 10^{-3}$ | 15 | *CFL1, PML, TGFB1* |
| | Positive regulation of muscle cell differentiation | $9 \times 10^{-3}$ | 6 | *EHD2, ENG, NIBAN2, TGFB1* |
| M.11 (299 genes) | Golgi membrane | $2 \times 10^{-3}$ | 30 | *ASAP2, MAN1A1* |
| | Regulation of nervous system development | $8 \times 10^{-3}$ | 30 | *IQGAP1* |

### Higher expression of mitochondrial genes in G2 and G3 muscles consistent with higher proportion of slow myofibers

In the M.13 module, with higher expression in G2 (VL, VM, and RF) and G3 (GL), the mitochondrial-related genes were enriched (Table 1). Eighteen out of 122 genes enriched for mitochondria in this module were hub genes, highly interconnected genes in the module (Table 1). To assess a potential impact on mitochondrial metabolic processes, we mapped the 122 genes to mitochondrial pathways (Figure 6). The most enriched processes were the respiratory electron transport chain in oxidative phosphorylation, the tricarboxylic acid (TCA) cycle, and beta-oxidation. This observation suggests a higher oxidative metabolism in G2 and G3, which is consistent with a higher proportion of slow myofibers.

**Figure 6.**
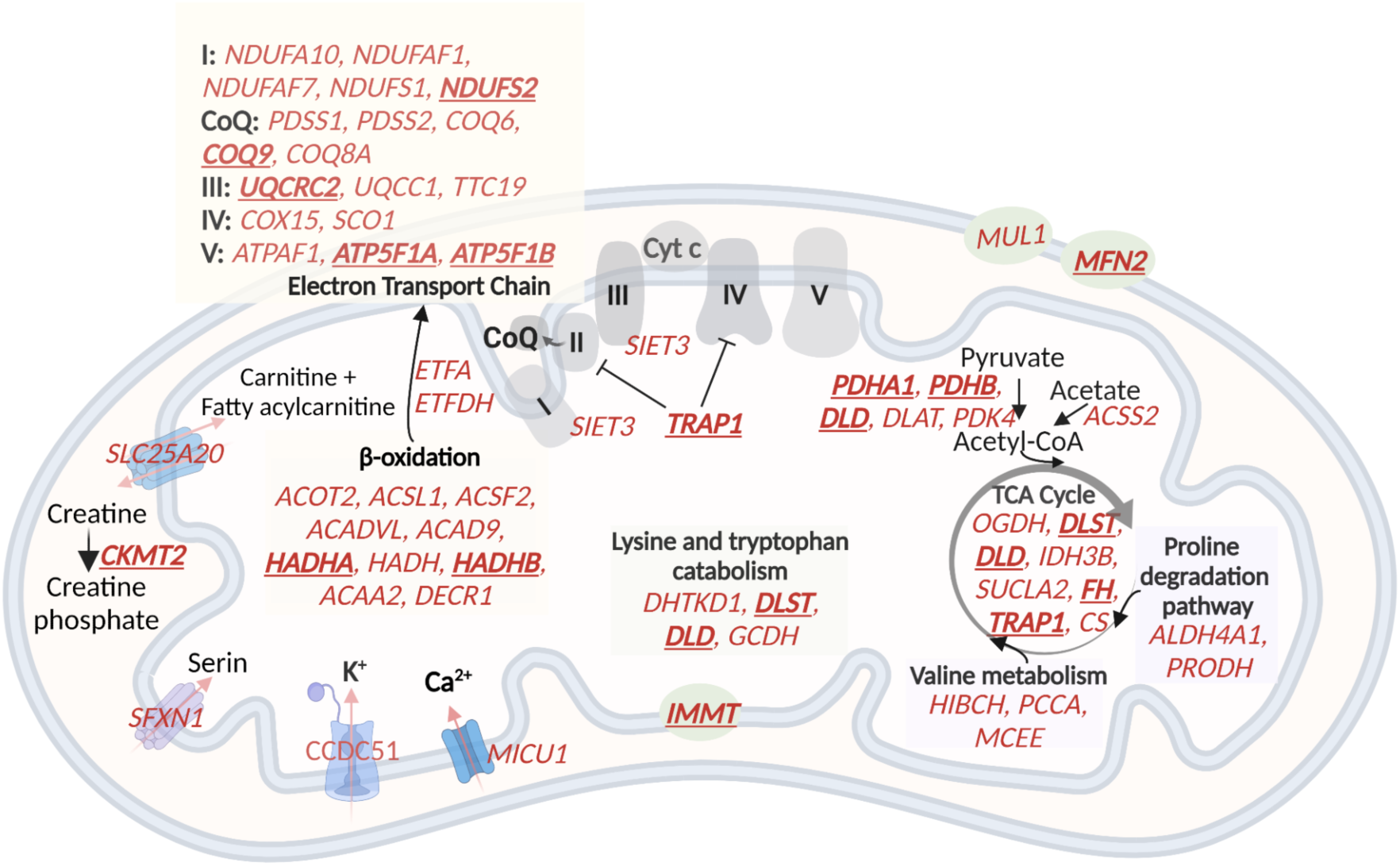
A schematic representation of genes with higher expression in G2 and G3, related to oxidative phosphorylation and metabolic pathways in the mitochondria. 60 (out of the 122) mitochondrial genes with higher expression in G2 and G3 are shown in red. The electron transport chain, lysin and tryptophan catabolism, TCA cycle, and beta-oxidation are shown. The hub genes are underlined and in bold.

### Homeobox transcription factors contribute to the mRNA diversity between the three groups of muscles

An enrichment for “anterior/posterior pattern specification” in M.14 was observed, with higher expression in G2 and G3 (Table 1). This module included *HOX* hub genes (Figure 7A). To assess whether the diversity between the groups of muscles was associated with the pattern of *HOX* gene expression, we plotted the normalized expression of all expressed *HOX* genes across all samples (Figure 7B). Remarkably, clustering based on *HOX* gene expression clearly separated the G1 from the G2 and G3 muscles (Figure 7B). Moreover, eleven out of 36 *HOX* genes were assigned to three of the muscle-related modules (M.14, M.30, and M.32), which showed the largest differences between the three groups of muscles (Figure 7B). Two *HOX* genes were selected (*HOXA10* and *HOXC10*) to further confirm the differences in expression between muscles using the *in situ* hybridization (ISH) procedure (Figure 8A). We included samples from GL and STM showing the largest difference in *HOX* genes expression. The *HOX* signal was mainly localized in myofibers (Figure 8A). Per sample, the average number of foci per myofiber was calculated revealing a higher number of *HOXA10* and *HOXC10* single molecule RNAs in STM compared with GL (Figure 8B). The ISH results were consistent with the RNA-seq data (Figure 8B-C), further demonstrating the robustness of our RNA-seq data.

**Figure 7.**
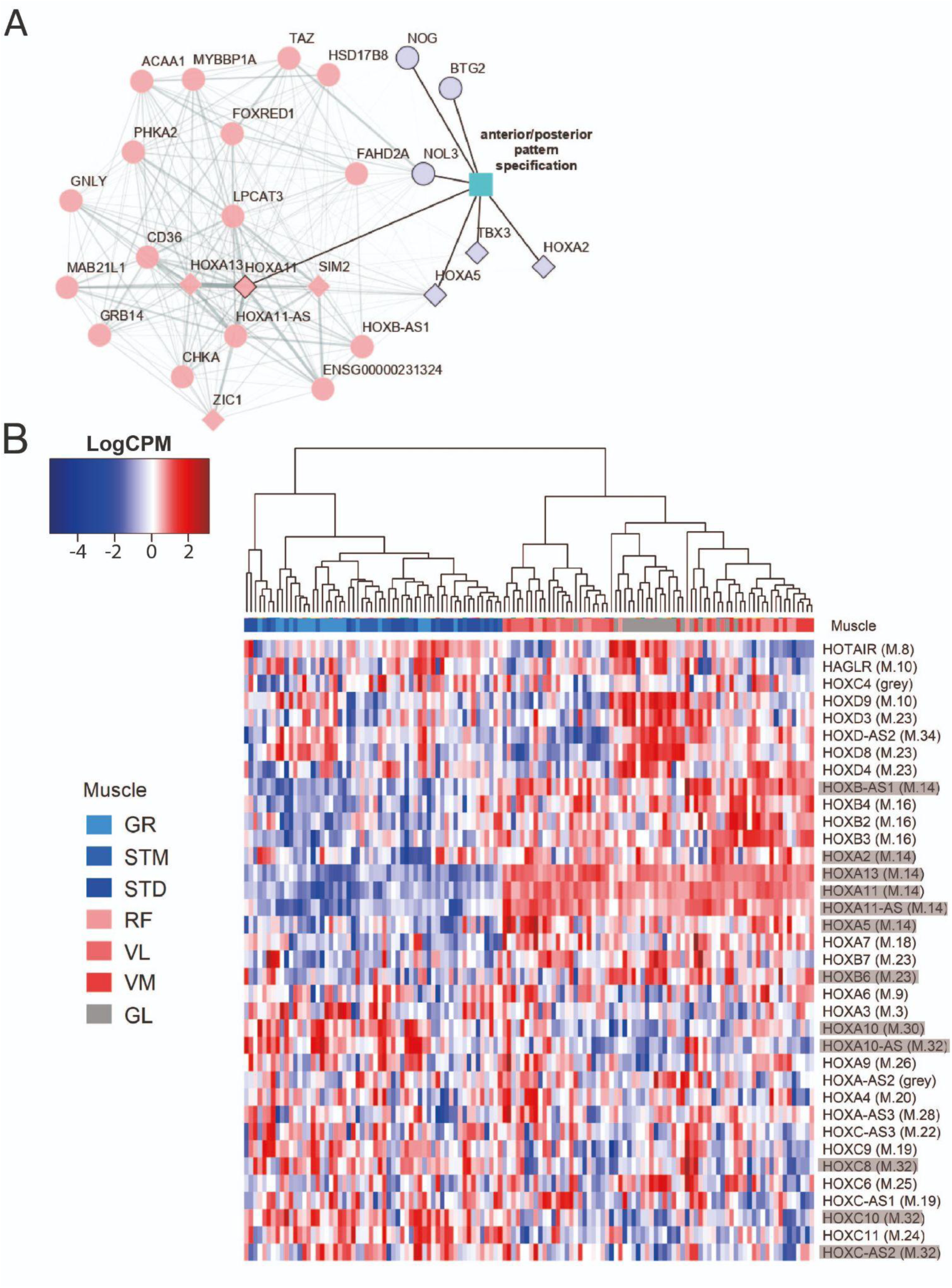
The expression patterns of *HOX* genes cluster muscles in the same groups. **A)** The graph shows the co-expression subnetwork of *HOX* genes and genes related to anterior/posterior pattern specification assigned to the M.14 module. Diamonds indicate transcription factors while other genes are indicated by circles. Pink and purple nodes represent the hub genes and non-hub genes, respectively. The genes related to anterior/posterior pattern specification have a black border. The edge thickness reflects the degree of topological overlap. Topological overlap is defined as a similarity measure between each pair of genes in relation to all other genes in the network. High topological overlaps indicate that genes share the same neighbors in the co-expression network. **B)** Normalized expression of all HOX genes (scaled by row) represented as a heatmap. The hierarchical clustering was generated using the normalized expression values. Each row represents a gene and each column represents a sample. The side color of columns indicates different muscles. The module in which the gene assigned is given between parentheses. Eleven highlighted HOX genes are assigned into muscle-related modules which showed the largest differences between the groups of muscles (M.14, M.30, and M.32).

**Figure 8.**
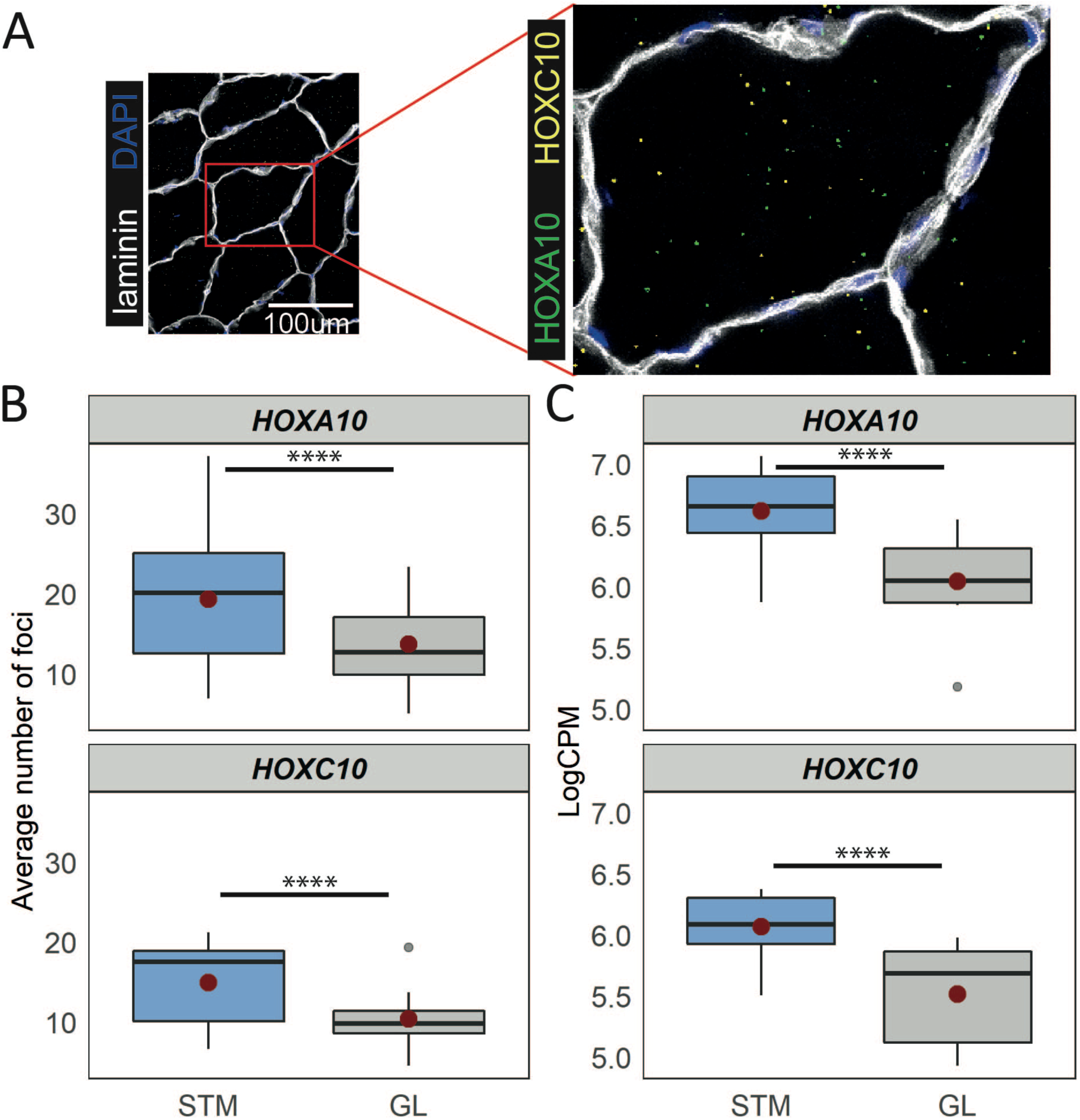
Distinct expression of *HOX* genes confirmed by RNAscope. **A)** A representative *in situ* hybridization image of *HOXC10* and *HOXA10* in muscle cryosections. The image is a merge image of the channels used for laminin, DAPI, *HOXC10* and *HOXA10* staining. **B)** The boxplots show the average number of foci per myofiber (y-axis) in STM and GL muscles (x-axis). **C)** The boxplots show the normalized expression of *HOXA10* (top) and *HOXC10* (bottom) in STM and GL muscles. The boxes reflect the median and interquartile range (N = 12 per muscle). The red dots on the boxes show the mean. **** *P-value* < 1×10^-6^ (linear mixed-model).

### Web application for exploring transcriptome atlas of human skeletal muscles

To facilitate data reuse and exploration of human skeletal muscle atlas, we developed a web application (https://tabbassidaloii.shinyapps.io/muscleAtlasShinyApp), enabling users to look up the sample information and the expression of any gene of interest. In addition, users can explore the list of genes used for the cell type composition analysis and their expression levels across all the samples. Furthermore, users can list and visualize the differentially expressed genes and the modules and their hub genes.

## Discussion

We generated a large skeletal muscle transcriptome atlas from 20 young healthy males. We included six leg muscles and two locations within one muscle. The atlas presented in this study is unique in terms of the number of muscles, the individuals included, and the age range of the participants. We confirmed the RNA-seq analysis using large-scale quantitative immunohistochemistry and mRNA in situ hybridization. Based on cell type composition, differential expression analysis and WGCNA, the seven leg muscle tissues consistently clustered into three groups: G1) GR, STM, and STD; G2) VL, VM, and RF; G3) GL. The muscles in G2 and G3 (VL, VM, RF and GL) showed higher proportions of slow myofiber types and higher capillary densities. GL, the only lower leg muscle, was distinct from VL, VM, and RF in its lower proportion of type 2A myofibers and a higher proportion of non-muscle cells. *HOXA10* and *HOXC10* expression were lower in VL, VM, RF and GL than in GR, STM, and STD muscles.

### Molecular diversity between muscles in different anatomical locations

The muscles included in this study mobilize and stabilize the knee joint. Muscles of the hamstrings (ST and GR) clustered together (G1), and muscles of the quadriceps (RF, VL and VM) clustered together (G2). The Hamstrings and quadriceps alternate in contraction and relaxation to flex, extend and stabilize the knee and aid in moving of the thigh. GL, the only lower leg muscle in our set, is located on the posterior side of the knee, allowing flexion of the knee and plantar flection of the ankle. Our study suggests that there is little molecular diversity between muscles of the same group, as compared to muscles in different groups of muscles.

We observed a higher proportion of fast-twitch myofibers in G1 compared with G2 and G3. This could be due to the role of the hamstrings in activities that require a large power output since fast-twitch myofibers are used more in these activities than slow-twitch myofibers (Bottinelli, Pellegrino et al. 1999, Willigenburg, McNally et al. 2014, Camic, Kovacs et al. 2015). Slow-twitch myofibers have a higher mitochondrial content compared with fast-twitch myofibers (Berchtold, Brinkmeier et al. 2000, Gouspillou, Sgarioto et al. 2014). Consistently, G2 and G3 muscles showed higher expression of genes encoding for mitochondrial proteins and a higher ratio of slow-twitch myofibers compared with G1. Slow-twitch muscles are also supplied by a denser capillary network (Nishiyama 1965, Murakami, Fujino et al. 2010, Korthuis 2011). Indeed, we observed a higher capillary density and higher endothelial cells in G3. The three groups of muscles also differed by the expression of *HOX* genes, specifically, *HOXA* and *HOXC* family members*. Hox* genes establish the anterior/posterior patterning during vertebrate embryonic limb development (Zakany and Duboule 2007). Interestingly, the development of these groups of leg muscles differs in developmental time (Diogo, Siomava et al. 2019), consistent with the expression of *Hox* genes (Zakany and Duboule 2007). *Hox* genes expression is not limited to embryonic development, but was found also in adult mouse muscles (Houghton and Rosenthal 1999, Yoshioka, Nagahisa et al. 2021), and *Hoxa10* gene was differentially expressed across adult limb mouse muscles (Yoshioka, Nagahisa et al. 2021). Moreover, Yoshioka, Nagahisa et al. (2021) demonstrated that *Hoxa10* expression in adult satellite cells affects muscle regeneration in mice. Here, we show that both *HOXA* and *HOXC* gene family are expressed in myofibers, and their expression levels differs between leg muscles. Yoshioka, Nagahisa et al. (2021) also showed expression of *HOX* genes in adult human muscle tissues. Yet, Terry, Zhang et al. (2018) concluded that the expression pattern of *Hox* genes in adult muscles is insufficient to explain the mRNA expression diversity in adult mouse skeletal muscles. Whether *HOX* genes are transcriptionally active in adult myofibers is a subject for future studies.

### Potential relevance to muscles disease and aging

In several muscle-related diseases like muscular dystrophies (MDs), muscle weakness and pathological features like replacement of muscle tissue with fat start in specific muscles and spreads to others as disease progresses (Emery 2002). This pattern differs between diseases, and the reason for the disease-specific involvement pattern is unknown. Exploring the molecular signatures that contribute to the differences between muscles may elucidate the pathophysiology of these diseases. For example, in Duchenne muscular dystrophy (DMD), which is caused by mutations in the *DMD* gene, the quadriceps is involved earlier, whereas the hamstring muscles are less involved, and the GR is spared (Wokke, van den Bergen et al. 2014, Hooijmans, Niks et al. 2017). The observed higher expression level of the *DMD* gene in ST and GR may be related to the late involvement in DMD patients during disease progression. Accessing the expression level of genes and implementing a quantitative approach (Veeger, van Zwet et al. 2021) to evaluate the association between leg muscle architectural characteristics and gene expression levels could be performed for other muscle diseases.

### Regional differences within muscles

The molecular and cellular differences between the samples from distal and middle locations of ST were larger than differences between ST and GR (Supplementary Figure S2). One module of co-expressed genes, M.21, showed a different expression pattern between STM and STD. This module was enriched for the cellular amino acid catabolic process and monocarboxylic acid catabolic process (Supplementary Table S5). While the distal side of the ST muscle has a rounded tendon, the differences between STM and STD cannot be explained by contamination of tendon tissue or closer proximity to the tendon, because we did not find a difference in the estimated tenocyte proportions between biopsies collected from the distal and middle parts of the muscle (Supplementary Figure S2). The myofiber composition was different between the distal and medial part of the ST muscle (Supplementary Figure S2A, Supplementary Figure S3). A divergent myofiber type composition of biopsies from superficial and deep areas of the same human muscle was reported by Johnson, Polgar et al. (1973) for the GL, RF, VL, VM, *adductor magnus*, *soleus*, and *tibialis anterior* muscles in the leg and thigh. Bindellini, Voortman et al. (2021) also reported different proportions of MyHC2A myofibers in distal and middle parts of *tibialis anterior* in mice.

### Interindividual differences were larger than differences between muscles

Despite the narrow age range and an inclusion of only one gender in our study, we observed that the percentage of variance explained by the individual surpassed the variance explained by the muscles (Supplementary Figure S5F). This is in agreement with findings from Kang, Kho et al. (2005). The inter-individual variations are possibly resulting from genetic and environmental (activity, exercise, diet, etc.) factors. To account for inter-individual variation, we included the individual as a random effect in the different analyses and constructed a consensus gene co-expression network by merging the co-expression networks separately constructed per individual. Only after properly accounting for interindividual differences, we could identify the intrinsic differences between leg muscles.

### Study limitations

Differences in cell type composition between muscles are best captured using single-cell sequencing. Previous single cell (De Micheli, Spector et al. 2020, Rubenstein, Smith et al. 2020, Xi, Langerman et al. 2020) and single nucleus (Orchard, Manickam et al. 2021, Perez, McGirr et al. 2021) studies reported the cellular composition of adult human muscles, where single nucleus profiling is preferred because myofibers cannot be dispersed into single cell suspensions. The high costs associated with single-cell technologies are currently prohibitive for performing large scale analyses of >100 samples such as performed in our study. Here, we evaluated differences in cellular composition by deconvolution of bulk RNA-seq based on marker genes reported in single-cell studies. This approach appeared to be suitable for analyzing differences in cellular composition between large sets of samples, as we observed good consistency with immunohistochemistry-based analyses of myofiber type and endothelial cell composition. A limitation of the deconvolution approach is, however, that this only captures cell types for which discriminative marker genes are available.

We further acknowledge that RNA expression levels do not necessarily match protein abundance in muscles and do not reflect post translational modifications (Greenbaum, Colangelo et al. 2003, Liu, Beyer et al. 2016). Although a protein atlas could relate to muscle cell function better than RNA expression profiles, generating a genome-wide proteome in skeletal muscles is challenging, as muscle proteomes are dominated by the high abundance of high molecular weight sarcomeric proteins, and capturing the low abundance proteins is challenging. Despite this limitation, we showed consistency between results obtained by mRNA expression profiling and immunohistochemical staining of the proteins that were in focus in our study.

In summary, we demonstrated divergent molecular and cellular compositions between skeletal muscles in different anatomically adjacent locations. Overall, the consistency of the gene expression patterns, and the results obtained from the immunohistochemistry and RNA in situ hybridization experiments indicates the high accuracy and reliability of the transcriptome atlas generated in this study. Therefore, this atlas provides a resource for exploring molecular characteristics of muscles and studying the association between molecular signatures, muscle (patho)physiology and biomechanics.

## Materials and methods

### 1. Subject characteristics and biopsy collection

Healthy male subjects (aged 18-32) undergoing surgery of the knee for anterior cruciate ligament (ACL) reconstruction using hamstring autografts were recruited from outpatient clinics of two hospitals: Erasmus Medical Center and Medisch Centrum Haaglanden. Inclusion criteria included age, sex, and the amount of routine exercise. Subjects eligible for reconstructive ACL surgery were mobile, had full range of knee motion, minimal to no knee swelling and had physiotherapy until the surgery.

A total of seven biopsies were taken from six different leg muscles (Figure 1A). To study molecular differences within the muscle, two biopsies from the middle and distal sides of the semitendinosus muscle (STM and STD, respectively) were collected. During the surgery, the tendons of the gracilis (GR) and semitendinosus muscles were used to reconstruct the ACL, and biopsies from these muscles were taken directly from the graft after harvesting the autografts at the beginning of the operation. After the ACL construction, biopsies from gastrocnemius lateralis (GL) rectus femoris (RF), vastus lateralis (VL), and vastus medialis (VM) muscles were taken by percutaneous biopsy (modified Bergstrom (Bergstrom 1975)) using a minimally invasive biopsy needle. All biopsies were immediately frozen in liquid nitrogen and were kept at -80°C.

The study was approved by the local Medical Ethical Review Board of The Hague Zuid-West and the Erasmus Medical Centre and conducted in accordance with the ethical standards stated in the 1964 Declaration of Helsinki and its later amendments (ABR number: NL54081.098.16). All subjects provided written informed consent prior to participation.

### 2. Sample processing, RNA isolation, and cDNA library preparation

Biopsies were cryosectioned for RNA isolation, immunofluorescence staining, and *in situ* hybridization. For each sample, three cryosections of 16 µm thick were collected onto SuperFrost slides (Thermo Fisher Scientific, 12372098) and stored at −20°C prior to staining. For *in situ* hybridization, the cryosections were mounted on SuperFrost Plus Adhesion slides (Thermo Fisher Scientific, 12625336) and stored at −80°C. For the RNA isolation, cryosections were transferred into MagNA lyser green beads tubes (Roche, 3358941001). Then, they were homogenized in QIAzol lysis reagent (Qiagen, 79306) using the MagNA Lyser. Subsequently, total RNA was purified with chloroform. For samples from a subset of individuals, RNA was precipitated with isopropyl alcohol (Supplementary Table S1). For the other samples total RNA was mixed with an equal volume of 70% ethanol and further purified with miRNeasy Mini Kit (217004, Qiagen) using the manufacturer’s protocol (Supplementary Table S1). To evaluate the effect of two different RNA isolation protocols, RNA from five GR samples were isolated with both protocols (Supplementary Figure S5A). For both protocols, DNA was removed using RNAse-free DNAse set (Qiagen, 79254) using the manufacturer’s protocol. RNA integrity was assessed with the Agilent 2100 Bioanalyzer using Eukaryote Total RNA Nano chips according to the manufacturer’s protocol (Agilent BioAnalyzer, 824.070.709) (Supplementary Table S1).

Poly(A) library preparation was performed in four batches each with 39 samples at Leiden Genome Technology Center (LGTC, the Netherlands). Information on the RNA isolation protocol and library preparation batches used for each sample can be found in Supplementary Table S1. Samples from different muscles and individuals were equally distributed in each library batch to minimize a batch effect bias. Approximately 200ng of total RNA was used as starting material. mRNA was enriched using oligo dT beads (polyA+ bead-based enrichment), fragmented, and converted to cDNA using random hexamers and SuperScript III (Invitrogen). End-repair, A-tailing, and adapter ligation were performed using NEBNext chemistry (New England Biolabs) and xGen dual index UMI adapters (Integrated DNA Technologies) according to the manufacturer’s protocol. Finally, USER digest (New England Biolabs) and 15 cycles of library amplification were performed. Libraries were purified with XP beads and analyzed for size and purity on a Bioanalyzer DNA HS chip (Agilent BioAnalyzer, 5067-1504).

### 3. Bulk RNA-sequencing and analysis

Illumina sequencing was performed by GenomeScan BV (Leiden, the Netherlands) on a Novaseq-6000 producing paired-end 2 × 150 bp reads. Fastq files were processed using the BioWDL pipeline for processing RNA-seq data (v3.0.0, https://zenodo.org/record/3713261#.X4GpD2MzYck) developed by the sequencing analysis support core (SASC) team at LUMC. The BioWDL pipeline performs FASTQ pre-processing, RNA-seq alignment, deduplication using unique molecular identifiers (UMIs), variant calling, and read quantification. FastQC (v0.11.7) (https://www.bioinformatics.babraham.ac.uk/projects/fastqc/) was used for checking raw read QC. Adapter clipping was performed using Cutadapt (v2.4) (Martin 2011) with default settings, followed by checking the QC using FastQC. RNA-Seq reads’ alignment was performed using STAR (v2.7.3a) (Dobin, Davis et al. 2013) against the GRCh38 reference genome. PCR duplications were removed based on UMIs using UMI-tools (v0.5.5) (Smith, Heger et al. 2017). Gene read quantification was performed using HTSeq-count (v0.11.2) (Anders, Pyl et al. 2015). Ensembl version 98 (http://sep2019.archive.ensembl.org/) was used for gene annotation. Samples with less than 5M reads assigned to annotated exons were re-sequenced or excluded from all downstream analyses. A SNP calling was performed using GATK4 (v4.1.0.0) (McKenna, Hanna et al. 2010). Possible sample swapping was checked using an SNP panel with 50 SNPs (Yousefi, Abbassi-Daloii et al. 2018). The similarity for calls of these SNPs showed that two samples in the same RNA isolation batch were swapped. We revised the labels of these two samples in our dataset for downstream analyses.

We performed all the analyses in RStudio Software (v1.3.959)(RStudio-Team 2020) using R Statistical Software (v4.0.2)(R-Core-Team 2020). Samples with more than 5M reads assigned to annotated exons were included in all downstream analyses (Supplementary Figure S5B). The HTSeq count table was used to create a DGEList object using the edgeR Bioconductor package (v3.30.3) (Robinson, McCarthy et al. 2010). The filterByExpr function from the edgeR Bioconductor package was used to keep genes with 10 or more reads in at least 16 samples (the number of samples in the smallest muscle group). The dataset was normalized using the calcNormFactors function (considering trimmed mean of M-values (TMM) method) from the edgeR Bioconductor package.

### 4. Quality control and batch effect correction

We performed principal component analysis (PCA) to evaluate the main difference between samples in an unsupervised manner. Log-transformed expression values after normalization by counts per million (CPM) were used to calculate principal components using the base function prcomp with the center and scale argument set to TRUE.

We then performed the analysis of variance to determine the factors driving gene expression variations. We estimated the contribution of known biological (muscle tissues and Individuals) and technical (RNA isolation protocol, RIN score, initial RNA concentration, library preparation batch, sequencing lane, and library size) factors on variation of gene expression. Data transformed by the *voom* function from the limma Bioconductor package (v3.44.3) (Law, Chen et al. 2014, Ritchie, Phipson et al. 2015) was used to fit a linear model for each gene. We included all biological and technical factors as fixed effects and fitted the following linear model:

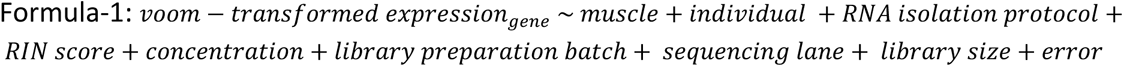

We used ANOVA from the car R package (v3.0-10) (Fox and Weisberg 2019) to estimate the relative contribution of each of these factors in the total variation of gene expression. Outcomes from both PCA and ANOVA revealed a strong library preparation batch effect (Supplementary Figure S5C and D), while the effect of other technical factors (RNA isolation protocol, initial RNA concentration, RIN score, and library size) was minimal (Supplementary Figure S5D). Accordingly, the HTSeq count table was corrected for the batch effect by the ComBat-seq Bioconductor package (Zhang, Parmigiani et al. 2020). The muscle was included in the ComBat-seq model to preserve possible molecular differences between muscles. The ComBat-Seq count table was used to create a DGEList object, followed by removing the low expressed genes and additional normalization using filterByExpr and calcNormFactors functions, respectively.

Outcomes of PCA and ANOVA confirmed the proper removal of the batch effect (Supplementary Figure S5E and F). In addition, the percentage of variance explained by the individual was found to be bigger than the variance explained by the muscle (Supplementary Figure S5F). We, therefore, included the individual as a random effect in all different analyses. Moreover, the RIN score was not considered as an exclusion criterion as it did not contribute to gene expression variation (Supplementary Figure S5F).

### 5. cell type composition estimation

We collected lists of genes marking different cell types that are present in human skeletal muscles from different studies (Smith, Meyer et al. 2013, Kendal, Layton et al. 2019, Perucca Orfei, Viganò et al. 2019, Rubenstein, Smith et al. 2020)(Supplementary Table S2). The expression of genes marking each cell type was summarized by their eigenvector (first principal component). We subsequently fitted a linear-mixed model to the eigenvector of each cell type using the lmer function from the lmerTest R package (3.1-3) (Kuznetsova, Brockhoff et al. 2017). These models included muscle as a fixed effect and individual as a random effect shown in the formula below:

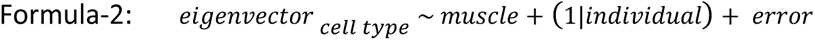

We tested the significance of fixed effects with the ANOVA from the car R package. The Benjamini-Hochberg false-discovery rate (FDR) was applied to adjust for multiple testing. We conducted post-hoc pairwise comparisons using the lsmeans R package (v2.30-0)(Lenth 2016) to identify a significant difference in the expression level of genes marking different cell types between different muscles. We used the pheatmap function from the pheatmap R package (v1.0.12) (https://CRAN.R-project.org/package=pheatmap) with the difficult setting to draw all the heatmaps.

### 6. Differential expression analysis (DEA)

We used the *voom*-transformed data to fit linear mixed-effects models for each gene using the lmer function from the lmerTest R package. The individual and muscle were included in the models as a random-effect and a fixed-effect, respectively, similarly to the formula-2. The *voom* precision weights showing the mean-variance trend for each observation were incorporated into the models. We tested the significance of fixed effects with the ANOVA from the car R package and the FDR was applied to adjust for multiple testing. We conducted post-hoc pairwise comparisons using the lsmeans R package to identify significant differences between each pair of muscles.

We calculated the Pearson correlation between differentially expressed genes (DEGs, FDR < 0.05) and the eigenvector of each cell type using the cor and cor.test from the stats R package. We adjusted for multiple testing using the FDR. DEGs which were significantly associated with a cell type eigenvector (Pearson correlation < 0.5 and FDR > 0.05) were defined as cell type related.

### 7. Consensus gene co-expression network analysis

In order to construct a gene network, we used the weighted gene co-expression network analysis algorithm using the WGCNA R package (v1.69) (Langfelder and Horvath 2008). We used the *voom* transformed data as an input. In order to calibrate the parameters of the network, we used the approach published by our group (Abbassi-Daloii, Kan et al. 2020). Briefly, prior knowledge of gene interactions from a pathway database was used to select the most optimal set of WGCNA parameters. We used the biweight midcorrelation (median-based) function in WGCNA of the signed hybrid type to define the adjacency matrix. We performed a full parameter sweep, testing various combinations of settings for power (6, 8, 10, 12, 14, 18, and 22), minClusterSize (15, 20, and 30), deepSplit (0, 2, and 4) and CutHeight (0.1, 0.15, 0.2, 0.25, and 0.3). These different settings were assessed using the knowledge network obtained from the Reactome database using g:ProfileR2 R package (v0.2.0) (Kolberg, Raudvere et al. 2020). All possible pairs of genes were assigned into four different groups: (1) in the same module and in the same pathway, (2) in the same module but not in the same pathway, (3) not in the same module but in the same pathway and (4) neither in the same module nor in the same pathway. The enrichment factor 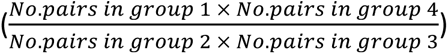 was calculated. The optimal set of parameters with the highest enrichment factor was: power: 8, MinModuleSize: 20, deepSplit: 0, Cut Height: 0.2. To identify gene co-expression networks that were consistent across individuals, we constructed first co-expression networks for each individual separately and merged these subsequently into a consensus co-expression network. To achieve this, the adjacency matrices per individual were raised to power 8 and converted into topological overlap matrices (TOM). TOM of some individuals may be overall lower or higher than TOM of other individuals. To account for this, we performed percentile (0.95) normalization over all the TOMs. The consensus TOM was then calculated by taking the elementwise 40th percentile of the TOMs. The consensus TOM was used to calculate the TOM dissimilarity matrix (*dissTOM* = 1 − *TOM*) which was then input to agglomerative hierarchical clustering (Langfelder and Horvath 2012). Finally, modules were identified using a dynamic tree-cutting algorithm from the resulting dendrogram (Langfelder, Zhang et al. 2008) specifying MinModuleSize = 20 and deepSplit = 0. The module labeled “grey” was not considered in the analysis as it consisted of genes that did not assign to any specific module. The summary expression measure for each module, the module eigengene (ME), was calculated (Zhang and Horvath 2005). Modules with similar expression profiles were merged at the threshold of 0.2. In addition, we calculated the intramodular connectivity to identify highly interconnected genes, called hub genes, per module.

### 7.1. Module-muscle association

To identify modules that differ in expression levels between muscles (named as muscle-related modules), we fitted linear mixed-effect models on the module eigengenes (MEs) using the lmer function from the lmerTest R package. These models included individual as a random-effect and muscle as a fixed-effect, like formula-2. We tested the significance of fixed effects with ANOVA from the car R package. We used ranova from the LmerTest R package to test the significance of random effects. To identify significant differences between each pair of muscles, we used a post-hoc multiple comparison tests as implemented in the lsmeans R package.

We performed a functional enrichment analysis for the muscle-related modules using ClueGO App (v2.5.7) (Bindea, Mlecnik et al. 2009) in Cytoscape (v3.8.1) (Kohl, Wiese et al. 2011). We used the CyREST API (Ono, Muetze et al. 2015) to execute the ClueGO by R script (http://www.ici.upmc.fr/cluego/cluegoDocumentation.shtml). Pathways and gene annotations from Kyoto Encyclopedia of Genes and Genomes (KEGG), Gene Ontology (GO), Reactome, and WikiPathways (WP) were included. The Benjamini-Hochberg FDR was applied to adjust for multiple testing. The annotations with any differentially expressed genes or hub genes or a transcription factor were included. To eliminate the redundant annotations, we only included an annotation with the lowest FDR for each ‘GoGroups’ defined by ClueGO and the annotations marked as ‘LeadingGoTerm’ by ClueGO.

We next determined muscle-related modules which showed the largest differences between the three groups of muscles. These modules were selected based on the FDR of GlueGO enrichment (< 0.01) and the F-value of the genes resulting from DEA in each module (third quantile > 5.5).

### 8. Immunofluorescence staining, imaging, image analysis

The immunofluorescence staining included myofiber typing and capillary staining. Prior to the staining, slides were allowed to equilibrate to room temperature, blocked for 30 minutes using 5% milk powder (FrieslandCampina, Amersfoort, The Netherlands) in phosphate-buffered saline containing 0.05% tween (PBST).

#### 8.1 Myofiber type composition

##### 8.1.1 Myosin staining

The antibodies for three myosin heavy chain (MyHC) isoforms (MyHC1, MyHC2A, and MyHC2X) and laminin were used as described by Riaz, Raz et al. (2016). Briefly, cryosections were stained with rabbit anti-laminin (1:1000, Sigma-Aldrich, L9393) and mouse anti-6H1 (1:5, DSHB; AB_2314830) detecting MyHC2X, for two hours at room temperature. Following the PBST washing, the secondary antibodies goat anti-rabbit-conjugated-Alexa Fluor® 750 (1:1000, Thermo Fisher Scientific, A21039) and goat anti-mouse-conjugated-Alexa Fluor® 488 (1:1000, A11001, Thermo Fisher Scientific) were incubated for an hour at room temperature. After PBST washing, sections were incubated overnight at four degrees with a mix of fluorescently conjugated monoclonal antibodies: BA-D5-conjugated-Alexa Fluor® 350 (1:600, DSHB, AB_2235587) and SC-71-conjugated-Alexa Fluor® 594 (1:700, DSHB, AB_2147165), detecting MyHC1 and MyHC2A, respectively. Lastly, after washing with PBST, the cryosections were mounted with ProLong™ Gold antifade reagent (P36930, Thermo Fisher Scientific) and stored at four degrees prior to imaging.

##### 8.1.2. Image acquisition, processing, and quantification

The stained slides were imaged with the Axio Scan.Z1 slidescanner (Carl Zeiss, Germany) using the ZEN Blue software (v2.6), capturing the entire section. The images were acquired with a 10×/0.45 Plan-Apochromat objective lens and the same image settings were used for all slides.

After imaging all cryosections, a shading profile was calculated using the ‘Shading Reference From Tile Image’ in ZEN Lite (v3.3) for each channel in each slide. This procedure produces a shading profile for each channel per slide and does not apply the shading correction. To improve the accuracy of the shading profile, we calculated the median over all the shading profiles over all scanned slides for each channel. These median shading profiles were then used to perform the shading correction using ‘Shading Correction’ in ZEN Lite (v3.3).

Further image processing was performed using Fiji (v 1.51) (Schindelin, Arganda-Carreras et al. 2012). Since the aggregated dataset is relatively large, we created a modular set of Fiji macros that process each step independently.

First, we converted the slidescanner datasets from the Carl Zeiss Image format (CZI) to multichannel 16 bit TIFF files using BioFormats (Linkert, Rueden et al. 2010). In this step, the images were 4x downsampled, by averaging, to improve the processing speed and reduce the dataset size. After downsampling the effective pixel size was 2.6 µm.

Next, we applied a semi-automated process to generate tissue masks from the laminin channel to determine the (parts of) cryosections to be quantified. To generate masks, we first used an automated procedure, inspired by ‘ArtefactDetectionOnLaminin’ method from MuscleJ (Mayeuf-Louchart, Hardy et al. 2018). Subsequently, a manual step was incorporated to check and correct the generated masks. For each sample, we performed manual corrections to remove artifacts such as tissue folds, out-of-focus regions, scratch, and dirt objects.

Then, we generated ‘masked’ copies of the laminin channel. To reduce any possible artifacts due to this binary mask, we applied a gaussian blur of 4 pixels to the masks and we set the pixel values of the laminin channel that were outside the mask to the median intensity of these pixels. The masked laminin images were then fed into the *Ilastik* pixel classification algorithm (Berg, Kutra et al. 2019). In *Ilastik* we trained a classifier to identify two classes: ‘myofiber boundary’ and ‘not myofiber boundary’. This classifier was then used to process all images in this dataset. This classification step greatly improved the subsequent laminin segmentation outputs.

Next, the laminin objects were segmented based on the output of the previous step. In short, the image was slightly blurred with a Gaussian Blur, after which the image was segmented using the Fiji method “Find Maxima” with output “Segmented Particles”, followed by binary dilation, and closing. Finally, the regions-of-interest (ROI) (individual laminin segmented objects) were generated using the ‘Analyze Particle’ method from Fiji.

After laminin segmentation, we measured the mean fluorescence intensity (MFI) as well as other properties in ROIs for all three channels using the Fiji measurement: “Mean gray value”. In addition, we recorded the “Area”, “Standard deviation”, “Modal gray value”, “Min & max gray value”, “Shape descriptors”, and “Median” features. We also quantified the results of the pixel-classification step by measuring its “Mean gray value” in each ROI as well as on the border (3-pixel enlargement) of each ROI. This quantification allows the assessment of the myofiber ‘segmentation certainty’, the certainty is high when the pixel-classification is high for the ‘myofiber boundary’ class all around the myofiber and low in the interior of the myofiber.

##### 8.1.3. Myofiber type composition analysis

First, we filtered out the non-myofiber objects since the laminin segmentation was automatic. We applied a percentile filtering for a ‘segmentation certainty’ on the cross-sectional area (CSA) and the circularity values. The objects with **(1)** pixel-classification on the object boundary less than 5^th^ percentile or **(2)** pixel-classification in the interior of the object greater than 95^th^ percentile or **(3)** CSA less than 10^th^ percentile or greater than 99^th^ percentile or **(4)** circularity greater than 1^st^ percentile were excluded. Samples from different muscles were pooled for all different filtering criteria except for the filtering for CSA, as the density distributions of CSA were found to differ between different muscles. In the next step, we selected the cryosection with the largest number of myofibers for each sample for further analysis. Samples with a minimum of a hundred myofibers were included in the myofiber type analysis. The final dataset contained 1,287,729 myofibers from 96 samples, with a median of 888 myofibers per sample. As previously described by Raz, van den Akker et al. (2020), per myofiber, the MFI values for each of three MyHC isoforms were scaled per sample (without centering). Subsequently, the composition of myofiber types was determined by clustering of the transformed (natural logarithm) MFI values. Each myofiber was assigned to a cluster using the mean-shift algorithm (bandwidth (*h*) = 0.02), a density-based clustering approach, implemented in the LPCM R package (v0.46-7) (Cheng 1995, Einbeck 2011). All the small clusters, with less than 1% from the total myofibers, were excluded. Then, per myofiber type cluster, the proportions of the total myofibers were calculated per sample.

#### 8.2 Capillary density

##### 8.2.1. Staining and image acquisition

Sections were stained with the primary antibodies: anti-human CD105 (endoglin, ENG) biotin-conjugated (1:100, BioLegend, 323214), anti-human CD31-Alexa Fluor® 594 conjugated (1:400, BioLegend, 303126), and rabbit anti-laminin for two hours at room temperature. After PBST washing, the slides were incubated with streptavidin-Alexa Fluor® 647 conjugated (1:500, Life Technologies, S21374) and goat anti-rabbit Alexa Fluor® 750-conjugated for an hour. After final PSBT washing, nuclei were counterstained with 4’,6-diamidino-2-phenylindole (DAPI) (0.5 μg/mL, Sigma-Aldrich) and were mounted with ProLong™ Gold antifade reagent. Cryosections were imaged with Axio Scan.Z1 slide scanner.

##### 8.2.2. Image processing and quantification

We used Fiji macros created for the myofiber composition analysis to convert CZI files to TIFF files, to generate the masks, and for the laminin segmentation. We then measured the cross-sectional area for laminin segmented objects using the Fiji “Area” measurement. Next, a Gaussian Blur filter with an σ value set to 1 was implemented on the CD31 channel, followed by thresholding using setAutoThreshold (“Li dark” algorithm) and processing using Watershed algorithm to separate touching and overlapping cells. The lumens were filled using the Fill Holes algorithm in Fiji. We then measured the properties in ROIs using the Fiji measurements: “Area”, “Mean gray value”, “Standard deviation”, and “Shape descriptors”. We then implemented the same processing on the ENG channel to select the ROIs but measured the “Mean gray value” and “Standard deviation in the CD31 channel to determine the CD31 and ENG colocalization.

For the image quantification, we first calculated the ratio between the total positively stained areas for CD31 and the total area of the muscle section, expressed as a percentage. We then determined the capillaries as the objects with (1) positive signals for both CD31 and ENG (Wehrhan, Stockmann et al. 2011), (2) larger than 3 µm^2^ and smaller than 51 µm^2^ (Poole, Copp et al. 2013), and (3) circularity larger than 0.5. Finally, we defined capillary density as the number of capillaries per unit (µm²) of muscle area.

### 9. RNAscope *in situ* hybridization

We detected single-molecule RNA using Multiplex Fluorescent Reagent Kit v2 (ACDBio, 323135) according to the manufacturer’s protocol for fresh-frozen cryosections, with the following adjustments to optimize the experiment for human muscles: fixation with 4% paraformaldehyde at 4 degrees for an hour, and all washing steps with washing buffer were performed three times for two minutes each. The protocol was optimized on control muscle cryosections by negative and positive probe sets provided by ACDBio. We performed the hybridization using probes for *Hs-HOXA11* (ACDBio, 1061891-C1), *Hs-HOXA10* (ACDBio, 867141-C2), and *Hs-HOXC10* (ACDBio, 803141-C3). Following the completion of the RNA probe hybridization, we carried out an immunostaining step at room temperature to label myofibers with rabbit anti-laminin followed by secondary labeling with goat anti-rabbit-conjugated-Alexa Fluor® 555 (1:1000, Abcam, ab150078). Lastly, following PBST washing, the nuclei were counterstained with DAPI (ACDbio, 323110). Cryosections were mounted with ProLong™ Gold antifade reagent. Slides were imaged with a Leica SP8 confocal microscope, equipped with a white light laser (WLL) source (Leica Microsystems, Germany) using a 40x/1.3 OIL objective. For each sample, multiple tiles at different regions across the muscle cryosection were images with seven z-planes (z-step size = 0.35 μm). The images for DAPI and *HOXA11* channels were acquired using a HyD 2 detector with 414nm-532nm excitation lasers and with 504nm-543nm excitation lasers, respectively. A HyD 4 detector was used to image anti-laminin and *HOXA10* channels with 558nm-585nm excitation lasers and with 603nm-665nm excitation lasers, respectively. A HyD 5 detector was used to image *HOXC10* channel with 675nm-800nm excitation lasers. The same image settings were used for all samples.

We performed the image processing in multiple steps and created a modular set of Fiji macros that process each step independently. We first merged and converted the Leica Image File (LIF) to a multichannel 16 bit TIFF file using the Grid/Collection Stitching Plugin (Preibisch, Saalfeld et al. 2009). We segmented myofibers using the following steps: 1) creating the maximum intensities projections of the laminin channel, 2) creating ‘probability’ maps of the laminin channel in *Ilastik*, 3) adding a point selection to the TIFF files, which seed the watershed, and 4) implementing watershed segmentation with two halting points for user interaction, first watershed segmentation and then making the ROI list (individual segmented myofibers) generated using the ‘Analyze Particle’ command.

After myofiber segmentation, we implemented a Gaussian Blur filter with an σ value set to 1 on each probe channel. We then applied the color threshold settings using setAutoThreshold (“RenyiEntropy dark” algorithm). Finally, for each probe channel, we measured the foci properties in each segmented myofiber using the Fiji measurements: “Area”, “Mean gray value”, “Standard deviation”, and “Shape descriptors”.

The RNA foci were defined as speckles smaller than 3.5 µm^2^ with circularity above 0.98. We excluded *HOXA11* from further analysis due to a low signal-to-noise ratio, agreeing with a lower expression level than *HOXA10* and *HOXC10*. Based on the negative controls, we defined threshold values to filter out false-positive signals for the 2 other HOX genes. These threshold values were set such that approximately all the foci in the negative control were classified as negative. Finally, to compare the expression of two genes between muscles, we calculated the average number of foci per myofiber per sample.

#### Availability of data and scripts

All scripts are publicly available on GitHub: github.com/tabbassidaloii/HumanMuscleTranscriptomeAtlasAnalyses. The raw data is publicly available at the European Genome Archive (Dataset ID: EGAS00001005904, https://ega-archive.org). Figure 1C and Supplementary Figure S1 show our analyses workflow used to explore genes contributing to the intrinsic differences between muscles.

#### Graphical user interface

The muscle transcriptomics atlas is available for exploration through a graphical user interface (https://tabbassidaloii.shinyapps.io/muscleAtlasShinyApp/) implemented using shiny, a web application framework for application shiny R package (v1.5.0)(Chang, Cheng et al. 2020).

#### Gene network visualization

The subnetwork was exported and visualized in Cytoscape (v3.8.1).

## Data availability

The raw data is publicly available at the European Genome Archive (Dataset ID: EGAS00001005904, https://ega-archive.org/). The muscle transcriptomics atlas is available for exploration through a graphical user interface (https://tabbassidaloii.shinyapps.io/muscleAtlasShinyApp/).

## Acknowledgments

We thank Susan Kloet, and the personnel from the Leiden genome technology center (LGTC) in the LUMC for providing the sequencing support.

## Supplementry data

### Supplementary Tables

**Supplementary Table S1.** Samples’ metadata.

**Supplementary Table S2.** A combined list of the genes marking different cell types.

**Supplementary Table S3.** The result of differential expression analysis.

**Supplementary Table S4.** List of genes – modules.

**Supplementary Table S5.** List of enriched biological processes and molecular functions within modules.

### Supplementary Figures

**Supplementary Figure S1.**
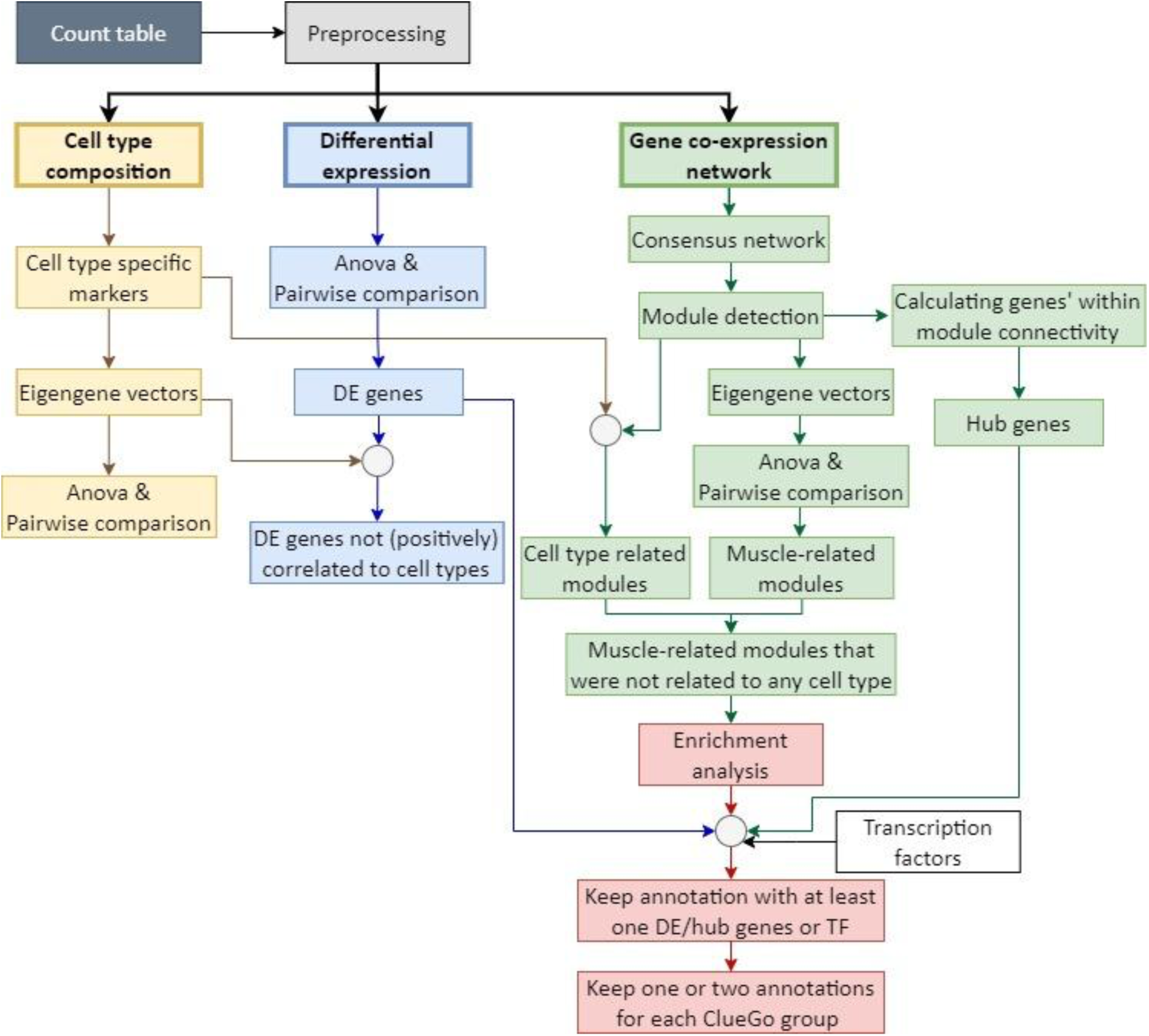
Analysis framework. A flowchart summarizing the analysis framework used to detect molecular signatures characterizing distinct skeletal muscles. Following pre-processing, muscle-specific signatures were identified using three approaches: cell type composition analysis (in yellow), differential expression analysis (in blue), gene co-expression network analysis (in green), and functional enrichment analysis (in red).

**Supplementary Figure S2.**
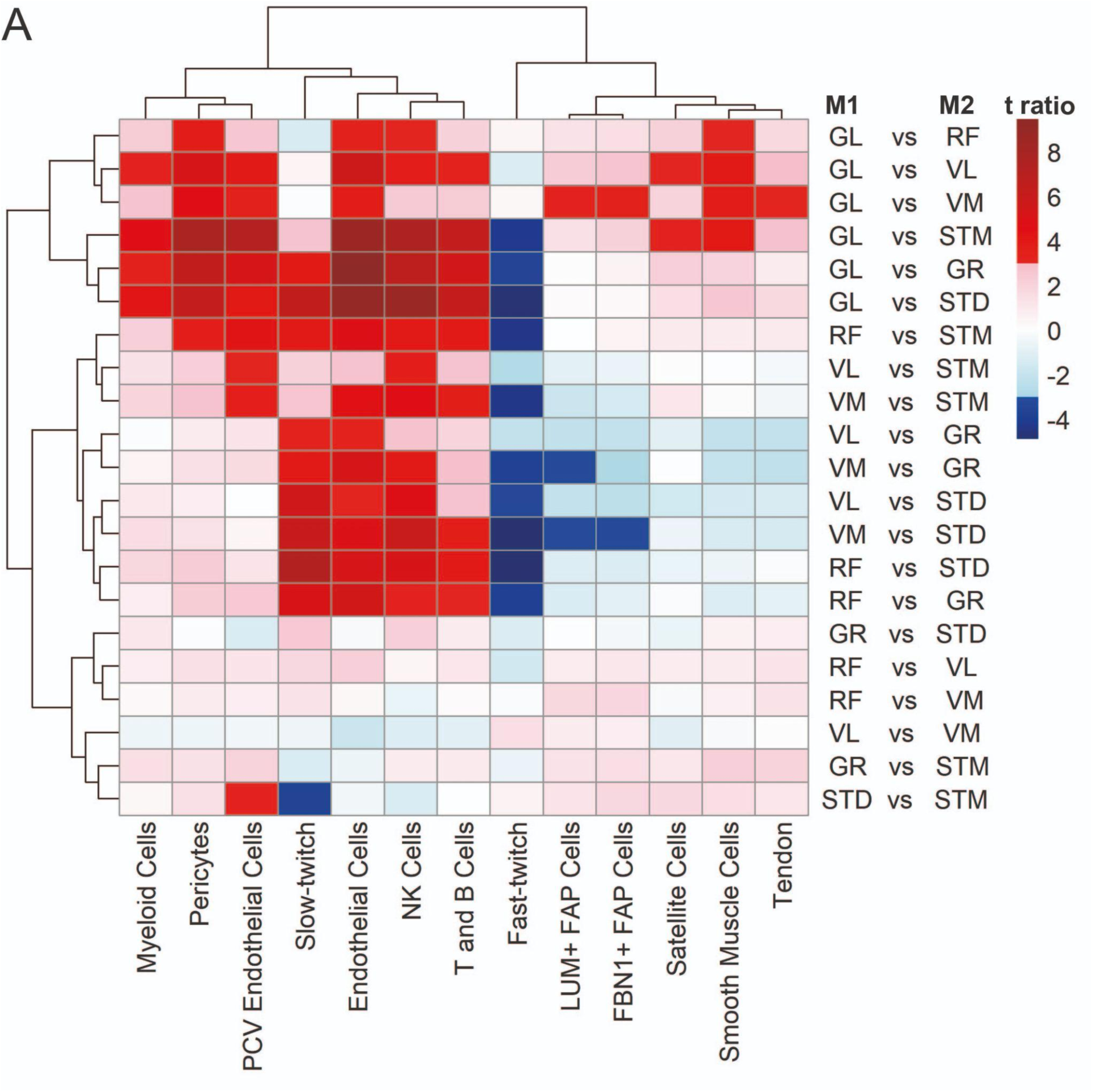
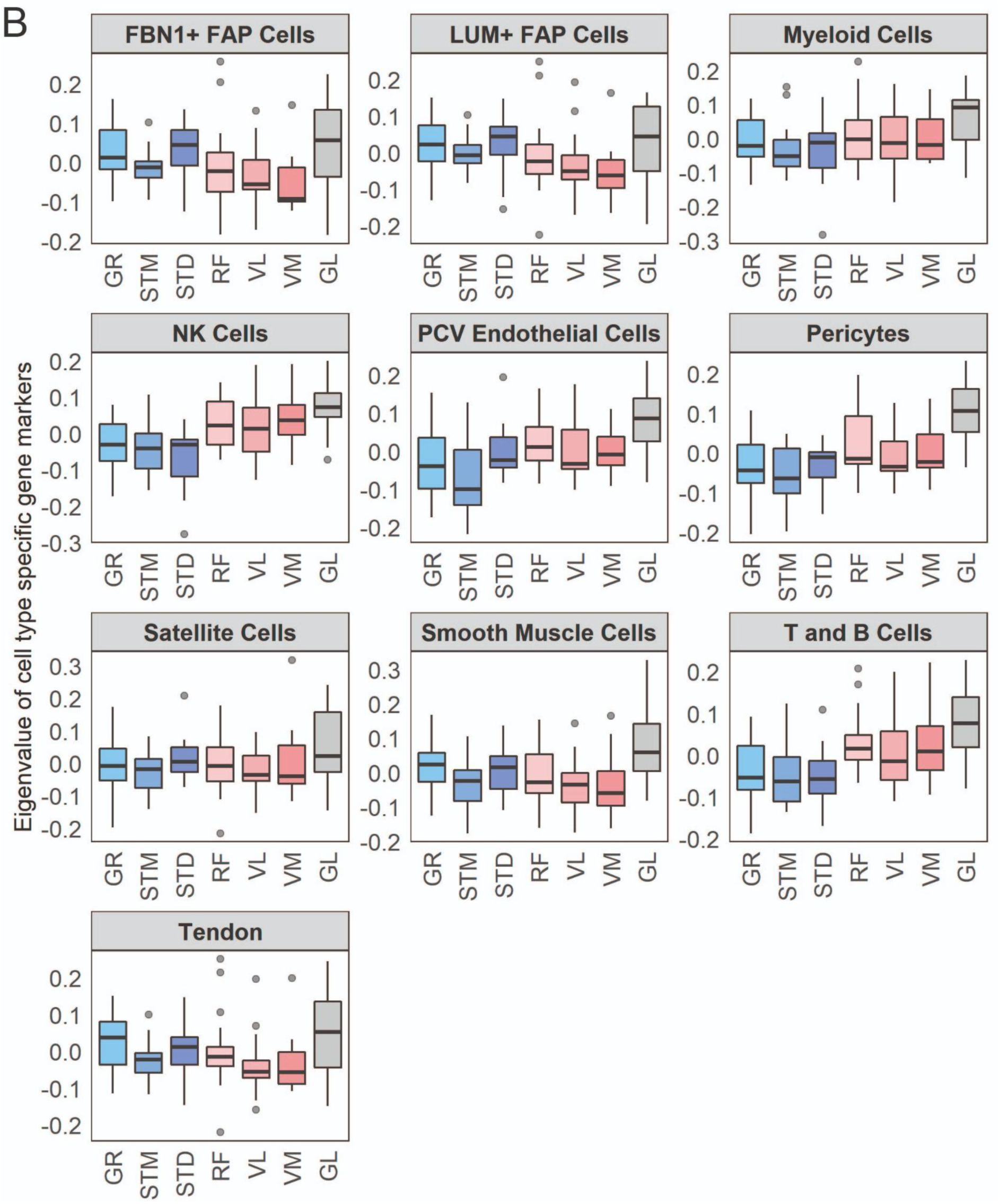
Cell type composition differences between muscles. **A)** The heatmap shows the differences between each pair of muscles. The statistically significant variation between muscles was tested by ANOVA followed by the post-hoc pairwise comparisons. Each row corresponds to a pairwise comparison, and each column shows a cell type. Color-coded cells show the corresponding t-ratio for the differences in eigenvalue of a cell type in each pairwise comparison. The significant differences (Tukey p-value < 0.05) are colored red (significantly higher eigenvalues in muscle M1) or blue (significantly higher eigenvalues in muscle M2). The non-significant differences are colored from pink (relatively higher eigenvalues in M1) to light blue (relatively higher eigenvalues in M1). **B)** Each boxplot shows the eigenvalues of a cell types across different muscles. The boxes reflect the median and interquartile range.

**Supplementary Figure S3.**
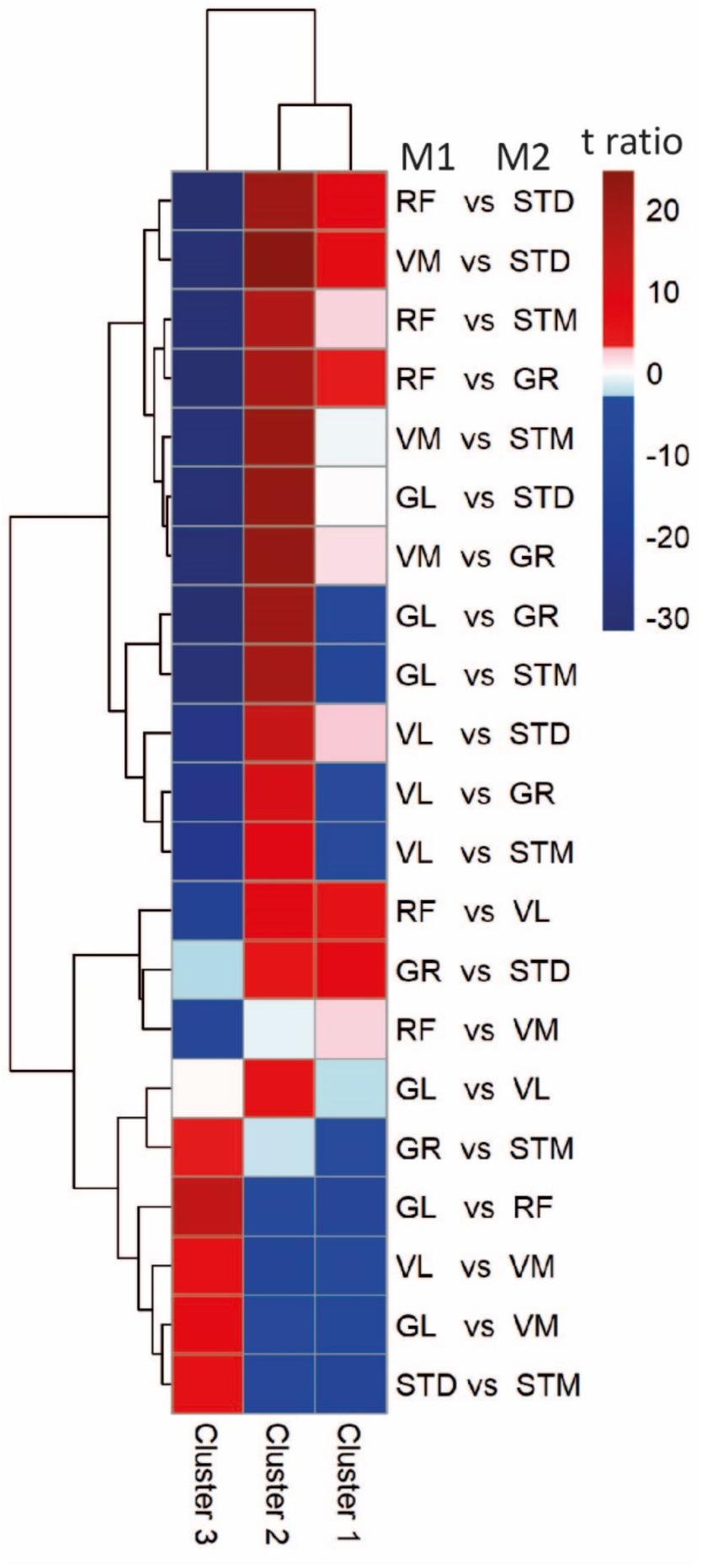
Myofiber composition differences between muscles. The heatmap shows the differences between each pair of muscles. The statistically significant variation between muscles was tested by ANOVA followed by the post-hoc pairwise comparisons. Each row corresponds to a myofiber cluster, and each column shows a pairwise comparison. Color-coded cells show the corresponding t-ratio for the differences in proportions of myofiber in each pairwise comparison. The significant differences (Tukey p-value < 0.05) are colored red (significantly higher proportions in muscle M1) or blue (significantly higher proportions in muscle M2). The non-significant differences are colored from pink (relatively higher proportions in M1) to light blue (relatively higher proportions in M1).

**Supplementary Figure S4.**
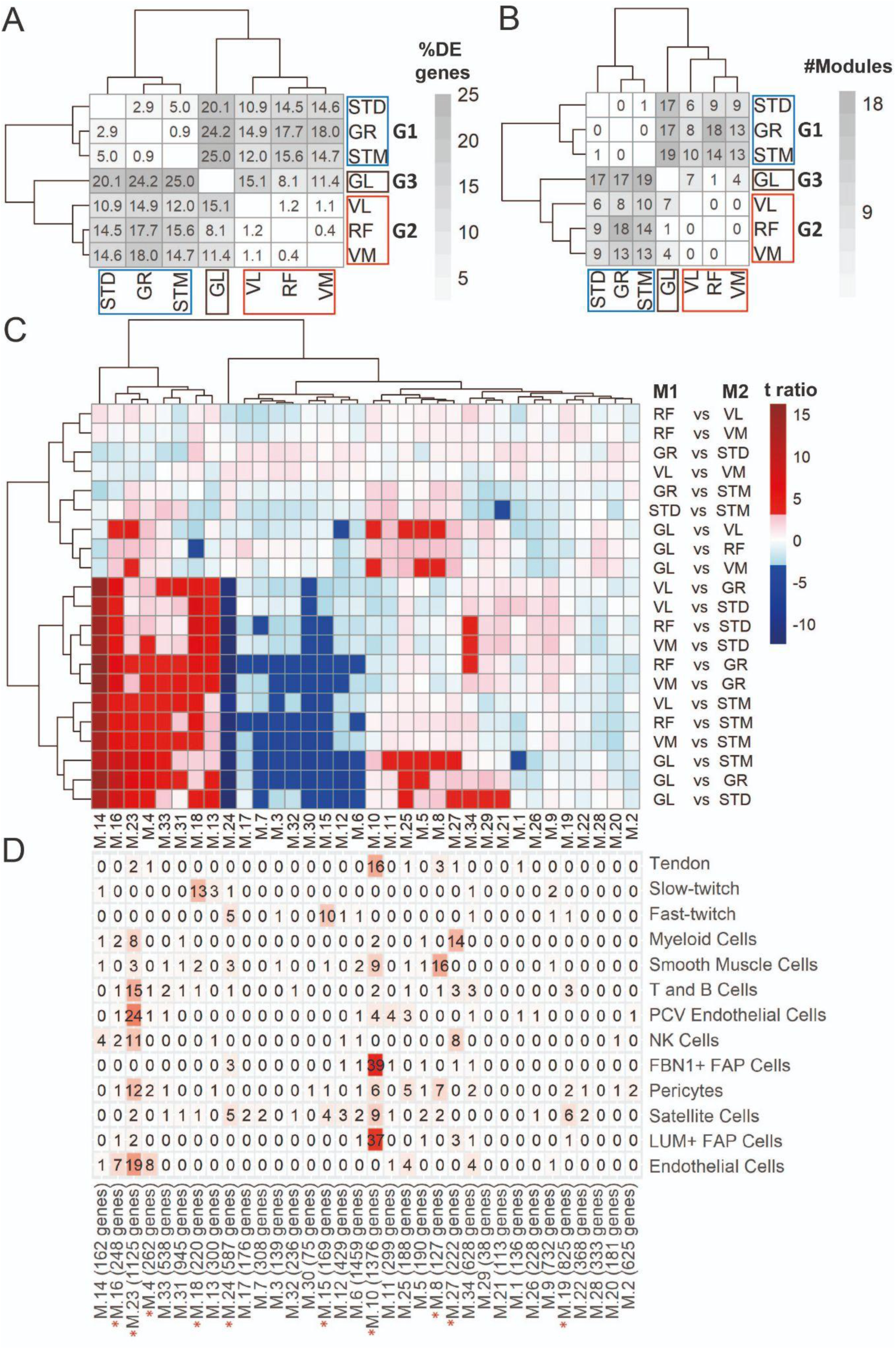
DEA and WGCNA also clustered muscles in three groups. **A)** Symmetric heatmap shows the proportion of all differentially expressed genes in different pairwise comparisons. Each row or column represents a muscle. **B)** Symmetric heatmap shows the number of modules that were significantly different in each pairwise comparison. Each row or column represents a muscle. **C)** Each row corresponds to a pairwise comparison and each column shows a muscle-related. Color-coded cells show the corresponding t-ratio for the differences in eigenvalue of a module in each pairwise comparison. The significant differences (Tukey p-value < 0.05) are colored red (significantly higher eigenvalues in M1) or blue (significantly higher eigenvalues in M2). The insignificant differences are colored from pink (relatively higher eigenvalues in M1) to light blue (relatively higher eigenvalues in M1). **D)** The table shows the intersection of the genes in the modules (columns) with genes marking different cell types (rows). Color-coded cells show the corresponding intersection number. The red asterisks show modules driven by cell type composition.

**Supplementary Figure S5.**
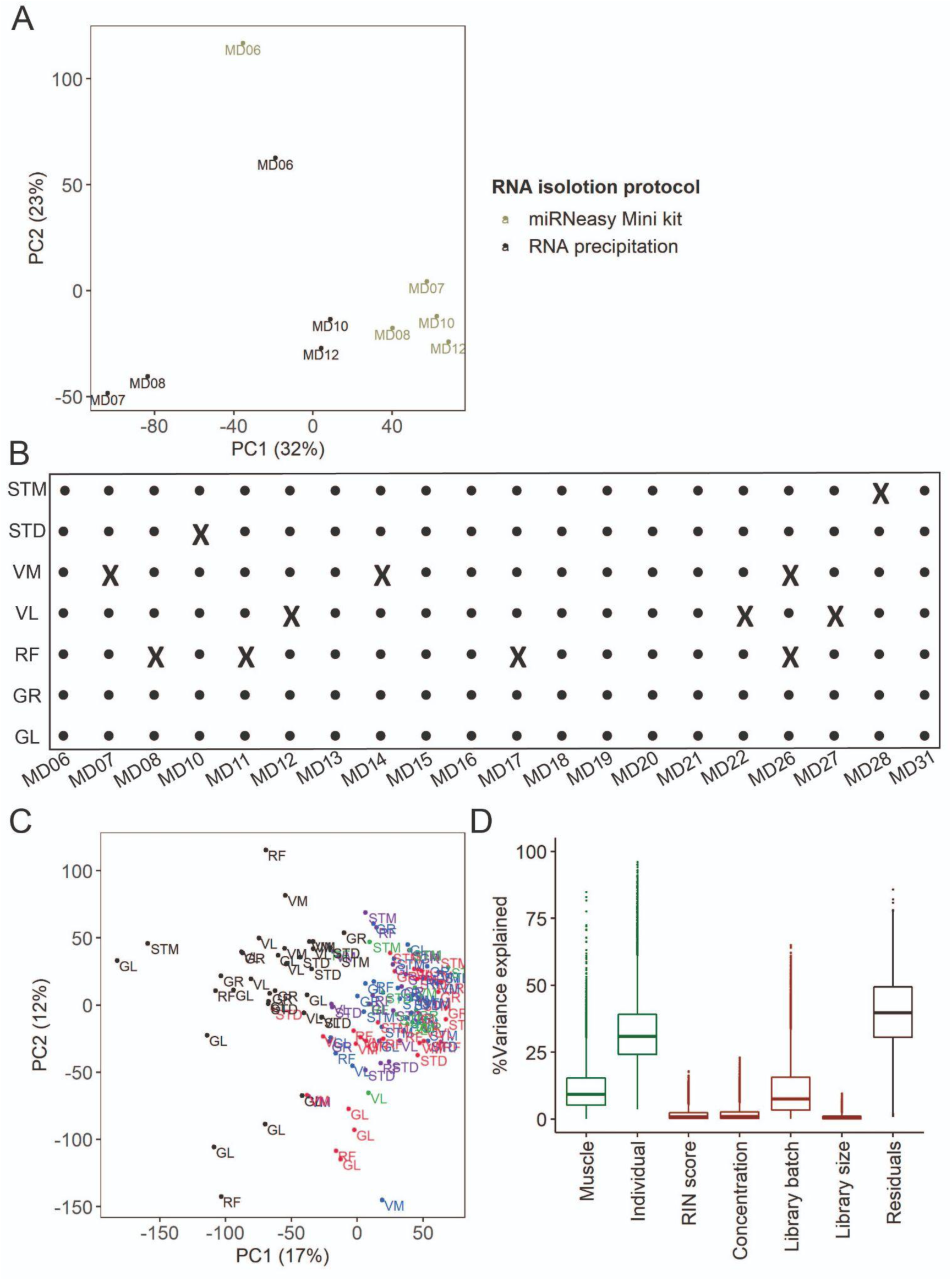
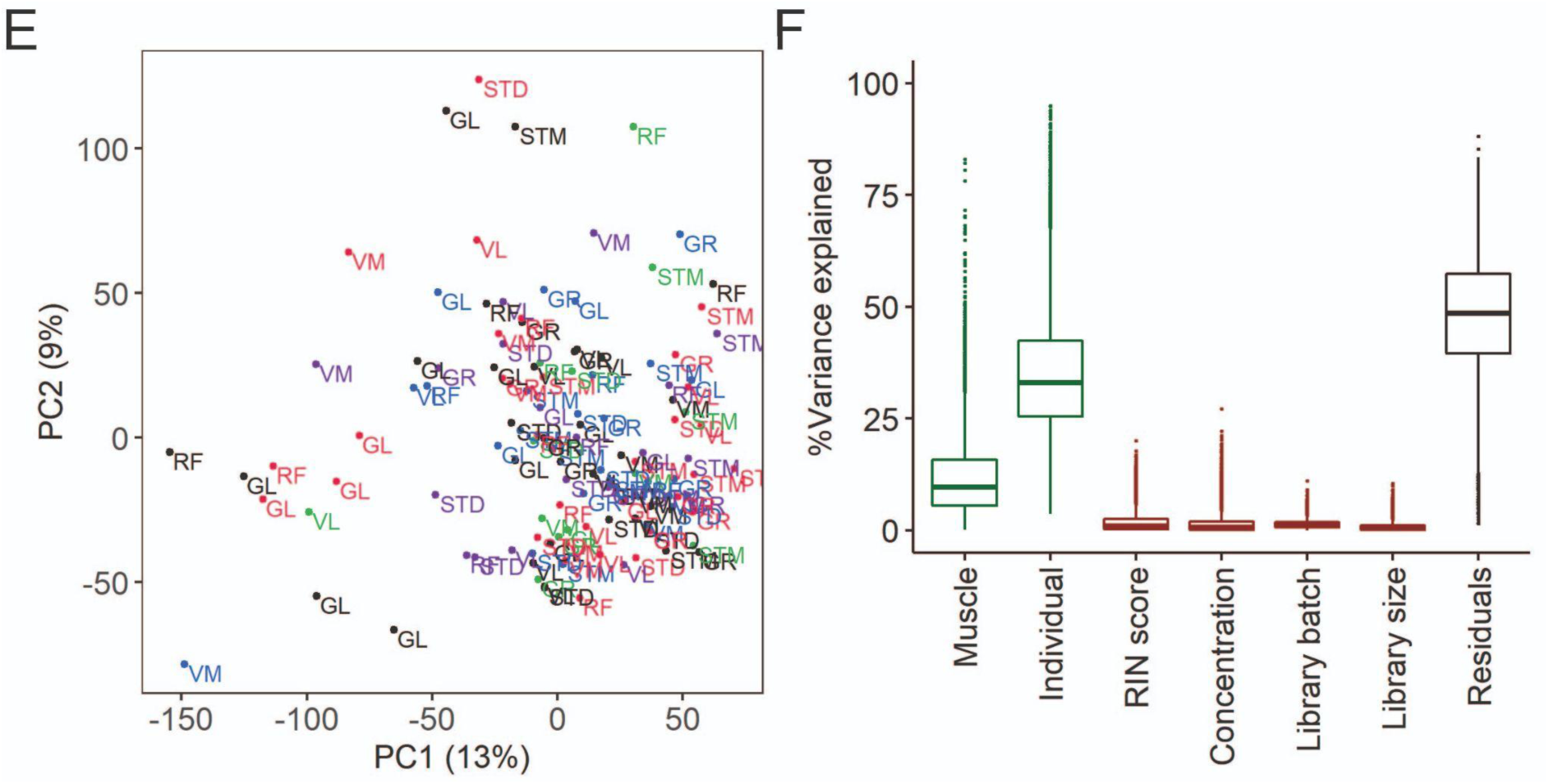
The quality control and batch correction of RNA-seq data. **A)** The PCA plot shows the effect of the RNA isolation protocols. The scatter plot of PC1 (x-axis) and PC2 (y-axis) shows GR muscle from five individuals from which RNA was isolated using two RNA isolation protocols. **B)** An overview of the RNA-seq samples (available from the European Genome Archive, Dataset ID: EGAS00001005904). An X indicates samples that are not present in the final transcriptome dataset. **C & E)** Scatter plots of PC1 (x-axis) and PC2 (y-axis) before (C) and after (E) batch correction. Each dot presents a sample labeled by muscle tissue. Each library preparation batch is shown with a different color. The re-sequenced batch is denoted in black**. D & F)** Box plots show the analysis of variance before (D) and after (F) batch correction. Y-axis shows the percentage of variance explained by different factors. The x-axis shows the known biological (muscle and individual, shown in green) and technical (RIN score, concentration, batch, library size, shown in red) factors. The RNA isolation protocol effect is captured in the individual effect.

